# A mechanism for sensing of and adaptation to K^+^ deprivation in plants

**DOI:** 10.1101/2020.03.21.000570

**Authors:** Feng-Liu Wang, Ya-Lan Tan, Lukas Wallrad, Xin-Qiao Du, Anna Eickelkamp, Zhi-Fang Wang, Ge-Feng He, Jian-Pu Han, Ina Schmitz-Thom, Wei-Hua Wu, Jörg Kudla, Yi Wang

## Abstract

Potassium ions (K^+^) are essential for manifold cellular processes. Organismal K^+^ homoeostasis requires sensing of K^+^ availability, efficient uptake and defined distribution. Roots are the organ for K^+^ uptake in plants and soil K^+^ availability shapes root growth and architecture^1^. Important channels and transporters conveying cellular K^+^ fluxes have been described^2,3^. Understanding K^+^ sensing and the mechanisms that orchestrate downstream responses exemplifies how environmental conditions integrate with root development and is essential to advance plant nutrition for sustainable agriculture. Here, we report where plants sense K^+^ deprivation and how this translates into spatially defined ROS signals to trigger HAK5 K^+^ uptake transporter induction and accelerated maturation of the Casparian strip (CS) paracellular barrier. We define the organ scale K^+^ pattern of roots and identify a postmeristematic K^+^-sensing niche (KSN) defined by rapid K^+^ decline and Ca^2+^ signals. We discover a Ca^2+^-triggered bifurcating low-K^+^ signalling (LKS) axis in that LK-enhanced CIF peptide signalling reinforces SGN3-LKS4/SGN1 receptor kinase complex activation. As consequence, activation of the NOXs RBOHC and RBOHD conveys transcriptome adaptation including HAK5 induction and accelerated CS maturation superimposed on the RBOHF-executed default CS formation. These mechanisms synchronise developmental differentiation and transcriptome reprogramming for maintaining K^+^ homoeostasis and optimising nutrient foraging by roots.

---

Roots confer K^+^ uptake from the soil using an elaborate array of transporters and channels to fortify this ion against a steep environment-organism gradient. While the cellular K^+^ concentration ranges around 100 mM, soils usually contain minute amounts in the sub-millimolar range. Soil K^+^ of 0.2 mM is sufficient to sustain normal growth of most plants, but lower supply provokes a suit of adaptive responses to protect plants against K^+^ deprivation^4^. A pivotal molecular facet of these responses is the switch from low affinity transport conferred by the channel AKT1 to high affinity uptake by the transporter HAK5 in Arabidopsis^5^-^10^. Moreover, enhanced accumulation of second messengers like Ca^2+^ and ROS has been associated with low-K^+^ (LK) responses^11,12^. On the organ scale, LK causes retardation/cessation of primary root growth, changes in root morphology (including promotion of lateral root and root hair growth), enhanced suberin shielding of the vascular stele and shut-down of root-to-shoot K^+^ transport^1,13-15^.

Primary root development involves controlled division and initial differentiation of stem cells in the meristematic zone (MZ), cell extension in the elongation zone (EZ) and subsequent cell file maturation in the differentiation zone (DZ)^16^. LK confers growth retardation by unknown LKS mechanisms through regulating stem cell activity via auxin distribution^15^. Lateral passage of K^+^ through the root towards the stele vasculature, which conducts ion transport to the shoot, involves flux across morphologically distinct cell files^17^. Of these, the endodermis surrounding the stele provides an inner border separating the stele long-distance transport compartment from outer cortex and epidermis cell layers^18^. Endodermis barrier function is brought about by genetically determined continuous maturation of the Casparian strip (CS) by lignin deposition and environmentally modulated further shielding by suberin deposition, which becomes enhanced upon K^+^ deprivation^19-23^. However, how K^+^ deprivation is sensed and signalled and how this connects with modulation of developmental programmes to bring about developmental plasticity remains to be elucidated. Here, we reveal where and how plants implement these responses to insufficient K^+^ supply.

## The root K^+^ landscape

We sought to determine the spatio-temporal pattern and dynamics of K^+^ concentration in the different cell types and tissues of Arabidopsis roots using the K^+^ reporter protein GEPII^24^. In roots grown with sufficient K^+^ supply (high-K^+^, HK) we discovered a surprising heterogenic distribution of K^+^ concentration. A sharp string of highest K^+^ concentration concurred with the mature vascular stele (Fig. 1). The outer edges of this K^+^ string were more diffuse only in the late elongation zone (EZ) and early differentiation zone (DZ). A second maximum of root K^+^ concentration occurred at the stem-cell niche (meristem). Radially, the different cell layers of the DZ exhibited unexpected layer-wise incremental inward increases of K^+^ concentration (Fig. 1e). K^+^ accumulation was barely detectable in epidermal cells including root hairs, intermediate in the cortex, further elevated in the endodermis and maximal in the stele. These findings suggest the existence and dominant function of multi-layered cell type-specific symplastic transport systems for organismic K^+^ acquisition and transport.

**Fig 1.**
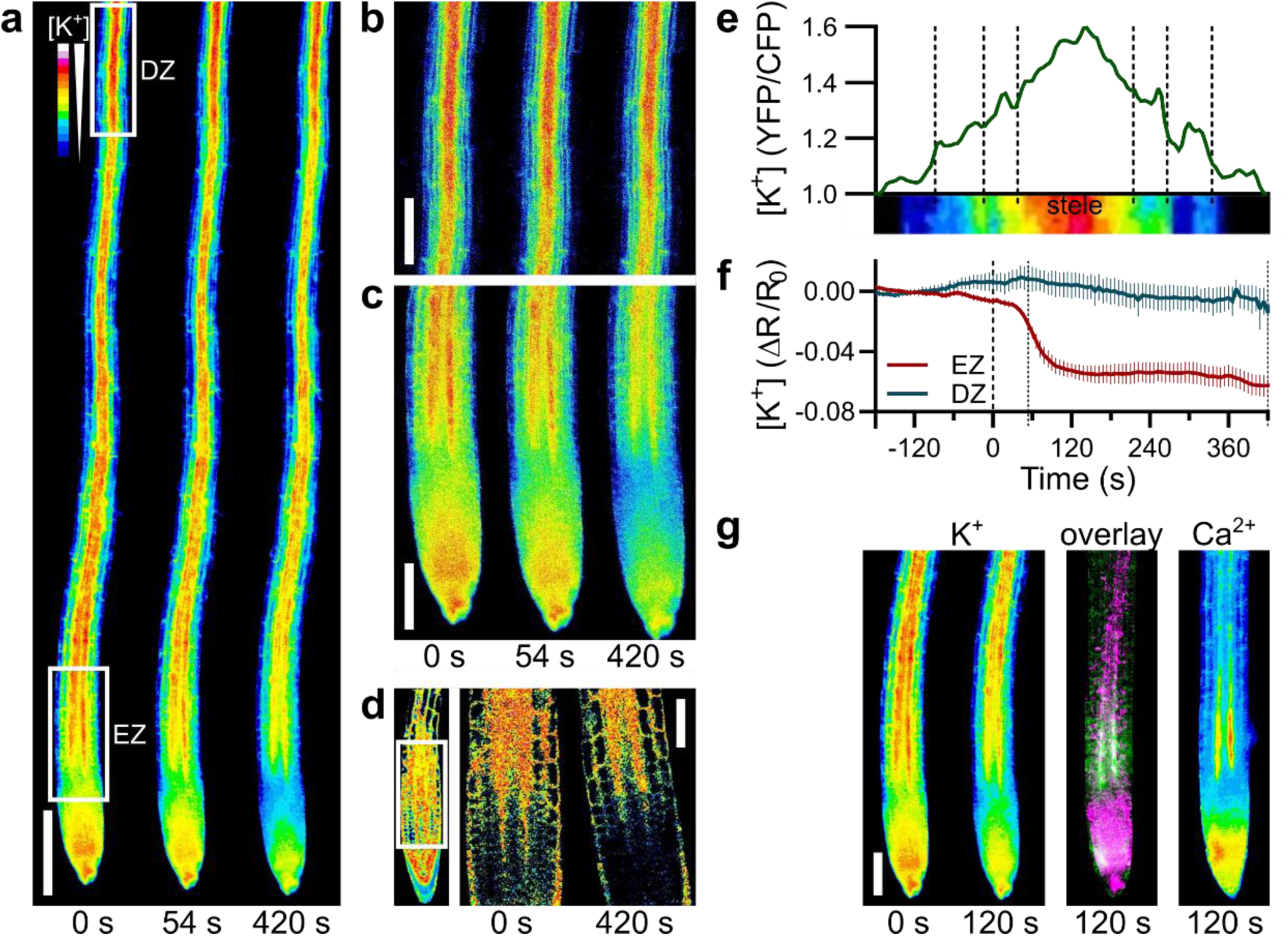
K^+^ imaging identifies cell-type specific K^+^ pattern and a K^+^-sensing niche. **a**, False colour representation of cytoplasmic K^+^ concentration ([K^+^]) at selected time points after onset of K^+^ depletion. Scale bar, 200 µm. **b** and **c**, Detail images of elongation zone (EZ) and differentiation zone (DZ) regions indicated in a. Scale bars, 100 µm. **d**, High-resolution imaging of [K^+^] in the K^+^-sensing niche (KSN). Scale bar, 50 µm. **e**, Transversal line-scan of K^+^ distribution in the mature (DZ). **f**, Quantitative determination of LK-induced cytoplasmic K^+^ dynamics in the (EZ) and DZ as depicted in a. Mean±SEM; n=5. **g**, False colour representation of K^+^ and Ca^2+^ accumulation in Col roots at indicated time points after onset of K^+^ depletion. The overlay displays the changes in K^+^ concentration between 0 s and 120 s in magenta and the Ca^2+^ signal after 120 s in green; white colour indicates the overlap of both signals.

We next assessed the impact of media K^+^ depletion on root K^+^ dynamics. Astonishingly, we observed an extremely site-restricted and rapid decline of K^+^ concentration initiating in less than 60 s that was strictly confined to a niche of postmeristematic cells at the MZ/EZ junction while no changes of K^+^ occurred elsewhere (Fig. 1 and Supplementary Video 1). This K^+^ decline became maximal within 120 s, stayed constant for at least 7 min (continuous measurement) and did not change during 48 h of low-K^+^ (LK) exposure (Fig. 1f and Extended Data Fig. 1a). Notably, in the remaining root the K^+^ concentration and pattern remained constant during this period. This niche-specific K^+^ decline temporally strictly coincided with the formation of a recently discovered LK-induced Ca^2+^ signal in these cells (Extended Data Fig. 1b)^11^. Moreover, spatially, the zones of K^+^ decline and Ca^2+^ elevation widely overlapped (Fig. 1g). Together, these data suggested that this dually responding postmeristematic cell group constitutes and functions as a K^+^-sensing niche (KSN) to orchestrate root signalling and adaptation to fluctuations in K^+^ supply.

## A peptide-activated receptor kinase complex conveys low-K^+^ signalling

Receptor complexes involving receptor-like kinases (RLKs) and associated kinases convey manifold sensing and signalling responses at the cell surface; yet their potential contributions to low-K^+^ signalling (LKS) remain elusive. We therefore screened a collection of about 200 RLK mutants by scoring leaf bleaching and growth retardation under LK and identified one mutant designated as *lks4-1* (*low-K*^*+*^ *sensitive 4*) as exhibiting respective phenotypes specifically upon K^+^ deprivation (Fig. 2a). *lks4-1* harboured a T-DNA insertion in the gene *At1g61590* encoding a receptor-like cytoplasmic kinase (RLCK) and causality of this mutation for the observed phenotypes was further corroborated by phenotyping of a second allele (*lks4-2*) generated by Crispr/Cas9 mutagenesis and by complementation analyses (Fig. 2a and Extended Data Fig. 2a, b). In both alleles, the K^+^ content of shoots was drastically reduced specifically under LK conditions while root K^+^ content was not significantly affected (Fig. 2b). Recently, *At1g61590* has also been described as Schengen1 (SGN1) that together with the RLK GSO1/SGN3, the NADPH oxidase (NOX) RBOHF and other proteins crucially functions in endodermis differentiation, Casparian strip (CS) formation and regulation of suberin sealing of this paracellular barrier^19,21-23,25-27^. We therefore addressed if the LK phenotypes of LKS4 would merely result from compromised barrier function or would indeed be indicative for impaired LK-signalling processes. To this end, we phenotyped various CS-related mutants. *sgn4/rbohF* did not exhibit K^+^ deficiency symptoms but in line with its endodermal defect displayed salt stress sensitivity as reported previously^25^ (Fig. 2c, d and Extended Data Fig. 2c, d). Moreover, we did not observe any discernible symptoms of K^+^ deficiency in mutants like *myb36, esb1* and *casp1 casp3* that all exhibit defects in CS positioning or formation (Fig. 2c, d and Extended Data Fig. 2c, d). In contrast mutation of either *SGN3* or *CIF2* rendered plants sensitive to LK and these mutants displayed reduced K^+^ accumulation similar to *lks4*. To investigate the mechanistic interconnection of LKS4 and SGN3 in LK-signalling we studied their potential direct interaction and phosphorylation. Bimolecular Fluorescence Complementation (BiFC) experiments indicated direct interaction of LKS4 with SGN3 at the plasma membrane but not with the related co-receptor kinase BAK1 (Extended Data Fig. 2e). In *in vitro* phosphorylation experiments SGN3 readily phosphorylated LKS4 but co-incubation of both kinases also caused enhanced phosphorylation of SGN3, suggesting that LKS4 may “feed-back” phosphorylate SGN3 or that direct interaction of both kinases further facilitates auto-phosphorylation of SGN3 (Extended Data Fig. 2f). These findings reveal a combined function of CIF2, SGN3 and LKS4 in the LKS pathway and that both kinases form an active receptor complex.

**Fig. 2.**
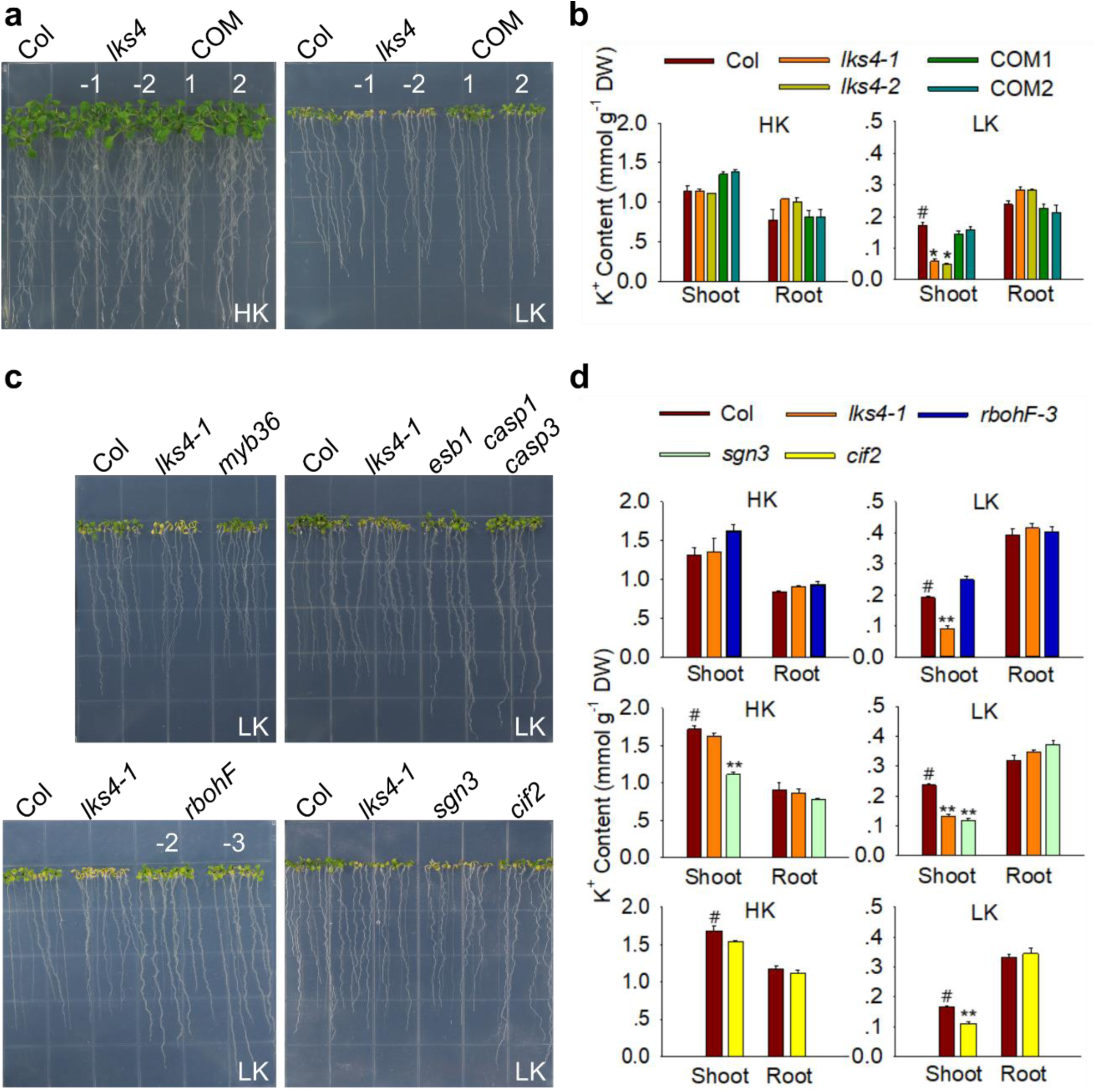
A peptide-activated receptor kinase complex determines plant K^+^ homeostasis. **a**, Phenotypes of WT (Col), two *lks4* alleles (*-1; -2*) and two complementation lines (COM) grown for 12 d on HK (high-K^+^=5 mM) and LK (low-K^+^=0.1 mM). **b**, Quantification of K^+^ content of plants depicted in a. Mean±SE; n=3; DW, dry weight; #, control; *p<0.05; **p<0.01. **c**, Phenotypes of the indicated lines on LK. **d**, K^+^ content of plants from d Mean±SE; n=3; DW, dry weight; #, control; *p<0.05; **p<0.01.

## An RLK-NOX axis for LK-induced ROS pattern

We next performed a phospho-proteomics approach comparing *lks4-1* and WT cultivated in either LK or HK to identify potential target proteins. This identified at least seven proteins including the NOX RBOHD as displaying a lower level of phosphorylation in *lks4-1* (Fig. 3a and Extended Data Tab. 1). We subsequently focused on RBOHD and other root-expressed NOXs as potential targets of LKS4 to mechanistically elucidate the contribution of ROS to LKS. To this end, we assessed the LK phenotypes of *rbohB, rbohC, rbohD* and *rbohF*. While *rbohB* and *rbohF* did not exhibit discernible LK phenotypes, *rbohC* and *rbohD* clearly exhibited LK-sensitivity and reduced shoot K^+^ accumulation although their symptoms were less pronounced than those of *lks4-1* (Fig. 3b, c and Extended Data Fig. 3a-c). The degree of suberisation in *rbohF* and *rbohD* was similar to that of WT arguing against differences in suberisation as the cause for the distinct LK-sensitivity of *rbohF* and *rbohD* (Extended Data Fig. 3d). We attempted to generate *rbohC rbohD* mutants but never obtained homozygous double mutants, suggesting an essential function of RBOHC/D-dependent ROS generation in plant development. To address genetically, if RBOHC and RBOHD function downstream of LKS4 in LKS, the respective double mutants were constructed and analysed. Both, *lks4-1 rbohC* and *lks4-1 rbohD*, displayed similarly sensitive phenotypes and a similar reduction of shoot K^+^ accumulation as *lks4-1* (Extended Data Fig. 3e-h).

**Fig. 3.**
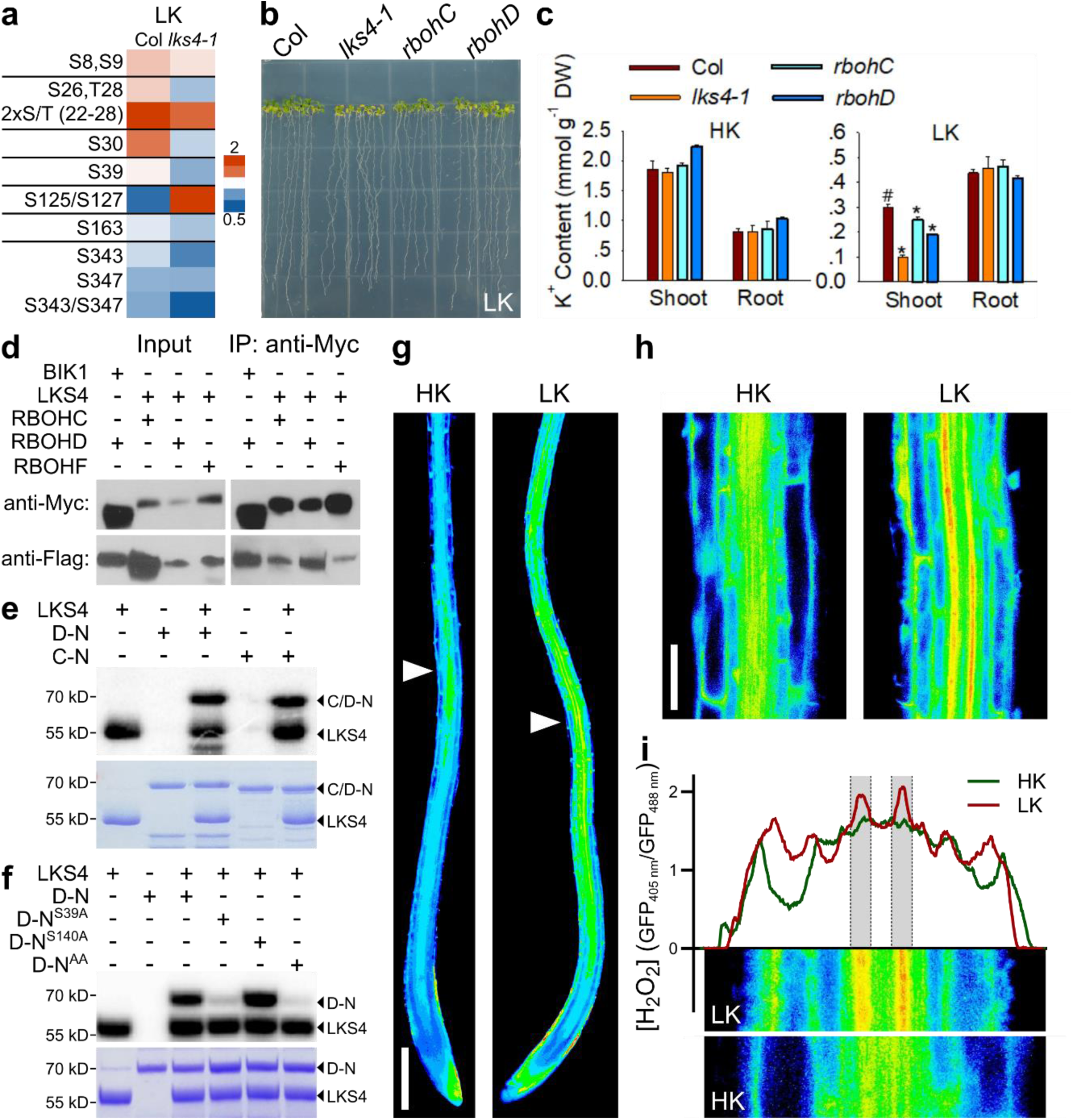
An RLK-NOX axis comprising RBOHC and RBOHD shapes LK-induced ROS pattern. **a**, Heatmap illustrating LK-induced differential phosphorylation of RBOHD in WT (Col) and *lks4*. 2xS/T (22-28) summarises two-site phosphorylation among S22/T24/S26/T28. **b**, Phenotypes of *rbohC* and *rbohD* grown on LK. **c**, Quantification of K^+^ content of plants depicted in b. Mean±SE; n=3; DW, dry weight; #, control; *p<0.05. **d**, NOX-LKS4 interaction analyses by Co-immunoprecipitation after expression in protoplasts. Proteins were detected with anti-Myc and anti-Flag antibodies before and with anti-Myc antibody after IP. **e**, Autoradiograph and Coomassie gel of *in vitro* phosphorylation assays combining LKS4/SGN1 with the N-termini of RBOHD or RBOHC. **f**, Autoradiograph and Coomassie gel of *in vitro* phosphorylation assays combining LKS4 with different phospho-site-mutated versions of the RBOHD N-terminus. **g**, False colour representation of cytoplasmic H_2_O_2_ accumulation after 24 h LK exposure. Arrows indicate root hair emergence. Scale bar, 200 µm. **h**, Detail images of regions indicated in g. Scale bar, 50 µm. **i**, Transversal line-scans of H_2_O_2_ accumulation. Arrows in g indicate positioning.

We next tested direct interaction of LKS4 with NOXs. In BiFC assays we reproduced the previously reported interaction of the RLCK BIK1 with RBOHD and detected interaction of LKS4 with RBOHC, RBOHD and RBOHF but not with RBOHB at the plasma membrane (Extended Data Fig. 4a). Furthermore, after co-expression of epitope-tagged versions of LKS4 with RBOHC, RBOHD or RBOHF in Arabidopsis protoplasts, we detected direct interaction of the kinase with all three NOXs (Fig. 3d). Phosphorylation assays revealed that LKS4 efficiently phosphorylated the N-termini of both RBOHC and RBOHD but did not phosphorylate their C-termini (Fig. 3e and Extended Data Fig. 4b). Together, these data suggest that direct LKS4-mediated phosphorylation of RBOHC and RBOHD can convey NOX activation for generation of LK-induced ROS signals and that the direct interaction of LKS4 with RBOHF may fulfil functions in other pathways, like CS-formation. Having established that LKS4 and RBOHC and RBOHD form a regulatory axis in LKS, we sought to delineate the mechanisms of LK-triggered NOX regulation. A combined *in vitro* phosphorylation/alanine-scanning approach identified S39 and S140 in RBOHD as phosphorylation sites targeted by LKS4 (Extended Data Fig. 5). While a S39A mutation largely reduced phosphorylation and a S140A mutation had only a minor effect, mutation of both serine residues almost completely abolished phosphorylation *in vitro* (Fig. 3f). To assess the functional and physiological significance of LKS4-mediated S39 phosphorylation *in planta*, RBOHD^S39D^ and RBOHD^S39A^ were expressed in *rbohD* and *lks4-1*. RBOHD^S39D^ fully rescued all LK-related phenotypes of *rbohD* and, remarkably, also significantly rescued these phenotypes in *lks4-1* (Extended Data Fig. 6a-e). Contrarily, plants expressing RBOHD^S39A^ did not exhibit phenotype complementation revealing the significance of RBOHD S39 phosphorylation by LKS4 for plant LKS.

We employed the ratiometric reporter protein roGFP2-Orp1 to investigate the spatio-temporal dynamics of LK-induced ROS generation in roots^28^. On HK, moderate H_2_O_2_ accumulation occurred in the basal DZ with maxima centred around the inner stele (Fig. 3g). Within 24 h of seedling transfer to LK, this pattern radically changed its dimension and intensity. While the lateral dimension of H_2_O_2_ pattern remained centred at the stele, its longitudinal dimension dramatically expanded to cover an area from the median EZ far into the DZ (Fig. 3g). High-resolution analyses of H_2_O_2_ pattern in the DZ revealed moderate H_2_O_2_ increases in all cell layers except the epidermis but dramatically increased H_2_O_2_ accumulation at the endodermis coinciding with the endodermis-specific expression of LKS4 (Fig. 3h, i). We next used the ROS indicator H_2_DCFDA to unravel the impact of the LKS4-RBOHC/D kinase-NOX axis on LK-induced ROS generation. On HK, a pattern of minor ROS accumulation similar to that observed with roGFP2-Orp1 was detected in WT and *lks4-1, sgn3, rbohC, rbohD* and in *lks4-1*/RBOHD^S39D^ plants (Extended Data Fig. 6f). K^+^ deprivation caused again intense ROS accumulation in WT that was strongly diminished in all mutants. *lks4-1*/RBOHD^S39D^ plants displayed a partially restored patchy ROS pattern (Extended Data Fig. 6g). Together these data establish the essential role of the SGN3/LKS4/RBOHC/D axis for LK-induced ROS signalling pattern in roots.

## ROS signalling triggers *HAK5* induction

The highly induced expression of the K^+^ transporter *HAK5* provides an adaptation mechanism to K^+^ deficiency and serves as molecular marker for LK responses^9^. We assayed the consequences of blocking LK-triggered ROS production on *HAK5* induction using RT-qPCR and *ProHAK5:LUC* lines. On HK, only minor basal luciferase activity was detected in WT, that was even lower in *lks4-1*. Exposure to LK strongly induced *ProHAK5*-driven luminescence in WT, but not in *lks4-1* and this LK-induction was completely blocked by the NOX inhibitor diphenylene iodonium chloride (DPI) (Fig. 4a). RT-qPCR revealed a clear dose-dependency of the DPI-mediated induction blockage (Fig. 4b). Upstream LKS pathway disruption in *sgn3* and *lks4-1* almost completely abolished LK-induction of *HAK5*, whereas *rbohC* and *rbohD* displayed less dramatic impairment of *HAK5* induction indicating their combined function in *HAK5* regulation (Fig. 4c). Expression of RBOHD^S39D^ in *lks4-1* partially restored *HAK5* induction. According with an essential upstream function of LKS4 for *HAK5* induction, leaf bleaching and K^+^ accumulation phenotypes of *hak5 lks4-1* were similar to those of *lks4-1*, while in *hak5* these phenotypes were less severe (Extended Data Fig. 7a, b).

**Fig. 4.**
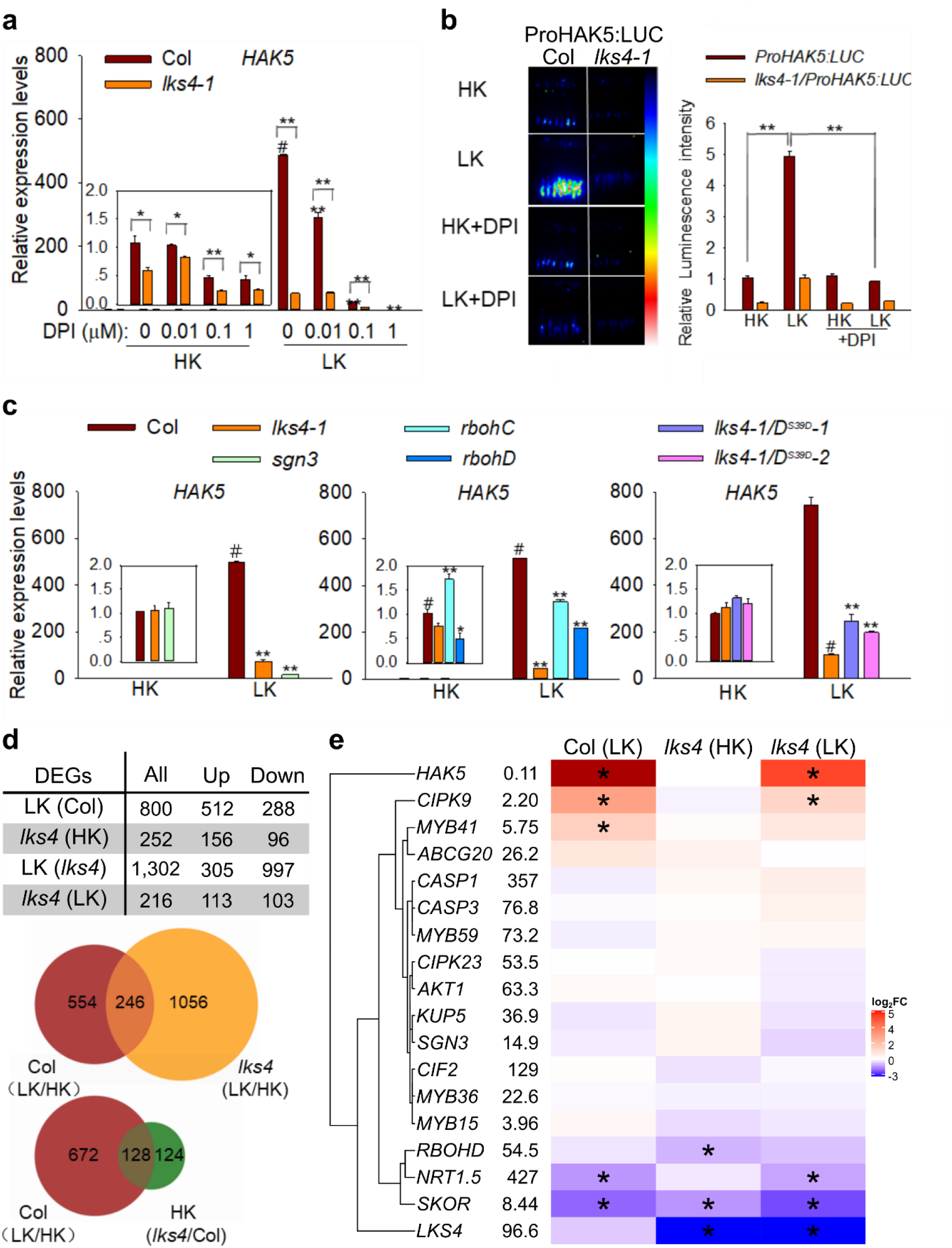
LK-triggered ROS generation is essential for *HAK5* expression induction and orchestrates specific LK-adaptive transcriptome reprograming. **a**, Impact of DPI treatment on the LK-induced expression of *HAK5* determined by RT-qPCR. **b**, Impact of DPI treatment on the LK-induced expression of *HAK5* determined by luminescence of a *ProHAK5:LUC* reporter. **c**, Determination of *HAK5* expression in the indicated genotypes by RT-qPCR. **d**, Summary of differentially expressed genes (DEGs) and Venn diagrams illustrating identified DEGs in Col and *lks4* upon exposure to HK or LK. **e**, Heatmap illustrating expression changes of genes related to K^+^ homeostasis and endodermal differentiation (*, DEGs with at least 2-fold expression change).

We next comprehensively defined the transcriptomic consequences of LKS and the role of LKS4 therein. In WT, LK exposure triggered differential transcript accumulation of 800 genes (512 upregulated, 288 downregulated) within 24 h (Fig. 4d). Remarkably, *HAK5* represented the far most induced gene exhibiting a 264-fold induction. Six further so far not experimentally characterised genes displayed a more than 150-fold induction (Supplementary Tab. 2a). Among enriched gene ontology (GO) terms we noticed cell-wall remodelling, oxidative stress, hydrogen peroxide catabolism, Ca^2+^ homeostasis, symporter activity, peroxidase and oxidoreductase activity reflecting the profound consequences of K^+^ deprivation on root growth and differentiation, ion transport and ROS generation (Extended Data Tab. 2). While the HK transcriptome of *lks4-1* differed in 252 genes from WT, LK triggered differential expression of 1302 genes in this mutant (Fig. 4d). Of these, only 246 genes were shared with the LK response of WT and only 79 genes were shared with the HK transcriptome in *lks4-1* (Fig. 4d and Extended Data Fig. 7c). In *lks4-1, HAK5* induction was reduced 12-fold down to 8 % of the upregulation in WT. Also, induction of 8 of the 10 most strongly induced genes in WT was largely diminished in *lks4-1* (Supplementary Tab. 2h). In contrast, the strong LK-triggered downregulation of *NRT1*.*5/NPF7*.*3* and *SKOR* (which both confer K^+^ loading into the xylem^14,29^) in WT was not exacerbated in *lks4-1* suggesting that the specificity of *LKS4* function allows for separating regulation of root K^+^ uptake from that of K^+^ loading into the xylem (Fig. 4e and Extended Data Fig. 7d). We also inspected the expression of genes related to endodermis modification including respective transcription factors (*MYB36, MYB15*), *CASP* genes, peroxidases and suberin related genes and did not detect noticeable expression changes for most of these genes (Fig. 4e and Extended Data Fig. 7e). Collectively, these results reveal that LK induces dramatic reprogramming of the root transcriptome and establish that LKS4 centrally functions in governing gene regulation in response to K^+^ deprivation.

## Bifurcating NOX branches in LKS

Since LKS4/SGN1 dually functions in endodermis modification/CS formation and in conferring LKS, we next scrutinised the prevailing paradigm that CS formation represents a “static” default developmental program not modulated by environmental cues. Functional CS barrier formation under LK and HK conditions was visualised using externally applied propidium iodide (PI) and revealed a significant 20 % accelerated CS formation in LK compared to HK (block of PI uptake; HK: 14±3 cells, LK: 11±2 cells after onset of elongation; Fig. 5a, c and Extended Data Fig. 8a). This surprising finding establishes that K^+^ availability modulates the developmental program of CS formation. Next, we sought to understand how LK triggers fortification of signalling via the CIF-RLK/RLCK-NOX axis. To this end, we first studied the impact of LK on CIF signalling peptide expression. This discovered a clearly discernible LK-triggered enhancement of CIF2 expression intensity in the DZ within 6 h that became full-blown after 24 h (Fig. 5b, d-e and Extended Data Fig. 8b). LK also accelerated the onset of CIF2 induction towards the meristem. The low expression level of CIF1 prevented reliable quantification of its expression (Extended Data Fig. 8b). Moreover, the expression pattern of LKS4 and SGN3 were shifted towards the meristem upon LK exposure (Extended Data Fig. 8c, d). These data reveal a molecular mechanism how LK triggers an accelerated and enhanced activation of the RLK/RLCK-NOX axis to stimulate CS formation. The co-occurrence of K^+^ decline and Ca^2+^ signals in the K^+^-sensing niche (KSN) raised the intriguing possibility that these Ca^2+^ signals may trigger upregulation of CIF2 for CIF-RLK/RLCK-NOX axis activation. Short-term application of 50 µM Ca^2+^ channel inhibitor LaCl_3_ (a concentration shown not to affect root growth and viability^11^) abolished the primary LK Ca^2+^ signal and completely abrogated the LK-induced changes in CIF2 expression pattern (Fig. 5b, d-e and Extended Data Fig. 8h). Accordingly, long-term LaCl_3_ treatment (50 µM) rendered WT plants sensitive to LK conditions and induced phenotypic alterations comparable to those of *lks4-1* challenged with LK in the absence of LaCl_3_ (Fig. 5f). These discoveries establish the absolute requirement of KSN-emanating Ca^2+^ signalling for activation of the CIF-RLK/RLCK-NOX axis to modulate root differentiation and establish LK resilience.

**Fig. 5.**
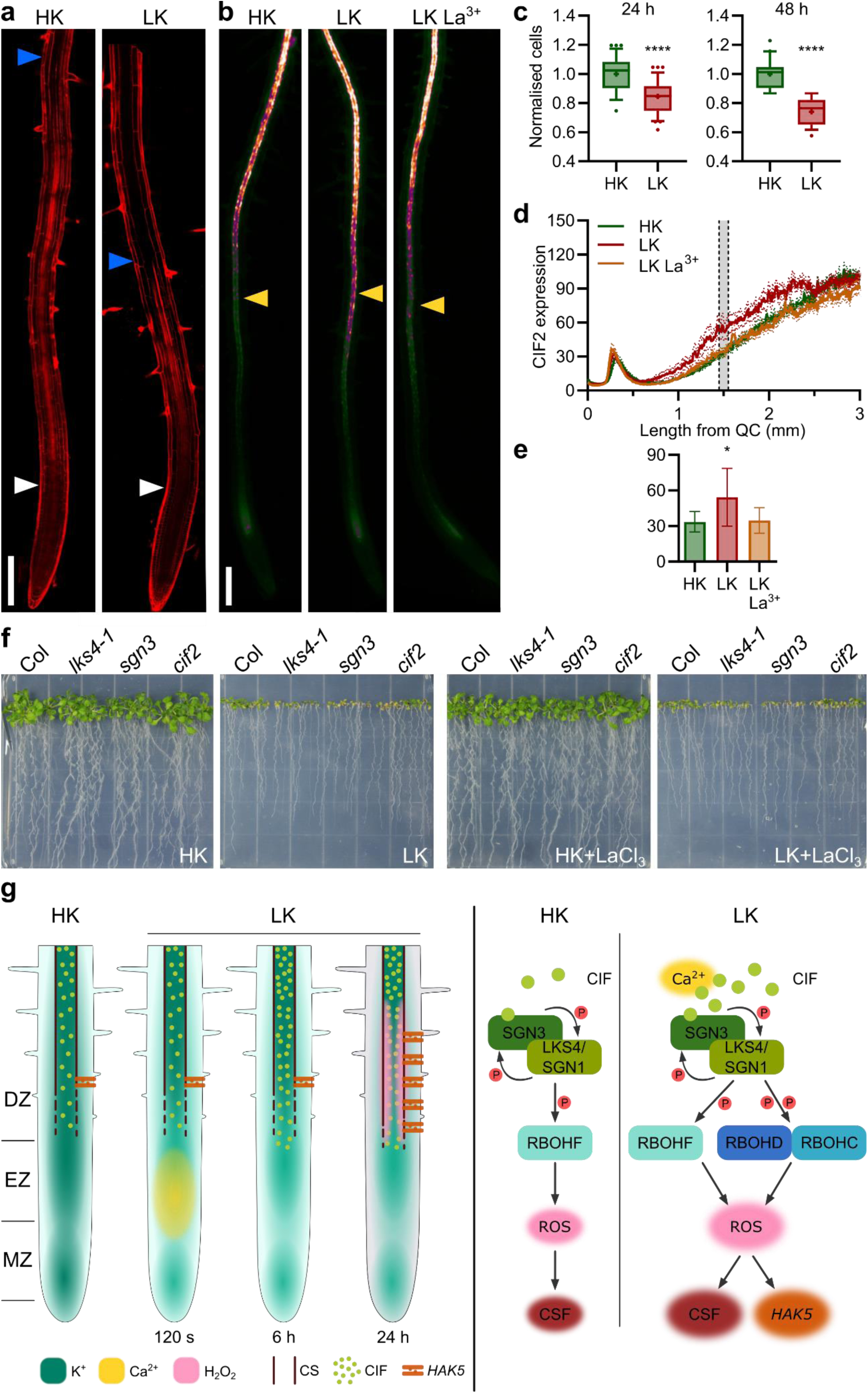
LK-induced Ca^2+^ signalling accelerates CIF peptide accumulation to impart LKS. **a**, PI-stained roots after 24 h HK or LK treatment. Arrows indicate the onset of elongation (white) and where staining of the stele ends (blue). Scale bar, 200 µm. **b**, CIF2 expression displayed as false colour GFP intensity of pCIF2:NLS-GFP expressing seedlings exposed to HK or LK for 24 h. La^3+^, incubation with 50 µM La^3+^ for 30 min prior to LK exposure. Scale bar, 200 µm. Yellow arrows mark the position 1.5 mm from the QC. **c**, Normalised cortex cell number from the onset of elongation until PI staining in the stele ends after 24 h and 48 h HK or LK exposure. Normalisation to the HK mean for each independent experiment. Median and 10-90 percentile; n=27, 34, 20, 21; HK 24 h, LK 24 h, HK 48 h, LK 48 h from 4 independent experiments; +, mean; ****p<0.0001; Two-tailed t test. **d**, Longitudinal quantification of CIF2 expression from b represented as 8-bit GFP intensity. Mean±SEM; n=9, 9, 11; HK, LK, LK La^3+^from 3 independent experiments. **e**, CIF2 expression at 1.5 mm from the QC. Mean±SD; *p<0.05; Dunnett’s multiple comparisons test). **f**, Phenotypes of indicated genotypes grown on HK±50 µM LaCl_3_ and LK±50 µM LaCl_3_ **g**, Proposed model for LK-sensing and adaptation in Arabidopsis. Left: Timeline of organ-scale events. The concentrations of K^+^, Ca^2+^ and H_2_O_2_ are illustrated by colour intensity. Interrupted lines indicate CS formation, while continuous lines indicate closed functional CS. Right, molecular model. CSF, CS formation, *HAK5*, transcriptome reprogramming.

Collectively, our findings allow to deduce a model of plant sensing and adaptation to K^+^ deprivation (Fig. 5g). Decline of soil K^+^ provokes an almost immediate drop in cytoplasmic K^+^ concentration in cells constituting the KSN. (The speed and cell-type specificity of this drop suggest active regulation as underlying mechanism instead of passive dilution.) Decreasing cellular K^+^ triggers plasma membrane hyperpolarization, possibly by activating proton (H^+^) ATPases enhancing H^+^ efflux. (Direct binding of K^+^ to the ATPase AHA2 inhibits its activity supporting a previously proposed ATPase activation by declining K^+^.)^30^ PM hyperpolarisation instantaneously triggers activation of Ca^2+^ channels to generate the LK-induced primary Ca^2+^ signal^31^. This accelerates and enhances CIF2 expression to strengthen SGN3/LKS4 receptor activation. This receptor reinforcement creates a bifurcating pathway for ROS generation, continuing to activate the NOX RBOHF for CS formation and inducing the additional activation of RBOHC and RBOHD for LKS. The activated LKS downstream pathway simultaneously involves accelerated CS formation and adequate transcriptome reprograming. Together, these synergistic mechanisms enable to enhance K^+^ uptake and to protect and sustain sufficient K^+^ concentration in the stele-born xylem to maintain organismic K^+^ homeostasis.

## Discussion

Here we report the organ scale landscape of K^+^ distribution and dynamics of Arabidopsis roots at cell/tissue resolution that profoundly advances current concepts of organismal K^+^ sensing and homoeostasis. We discover stepwise cell layer specific increases of K^+^ concentration that demand faithfully coordinated cell-type specific symplastic K^+^ fluxes. The resulting K^+^ maxima in the vascular stele remain preserved even under severe persistent deprivation of external K^+^ suggesting a physical necessity for upholding this high K^+^ value to allow for resuming root-to-shoot K^+^ fluxes upon resupply.

We identify the K^+^-sensing niche (KSN) formed by a group of meristematic/postmeristematic cells and defined by simultaneous rapid K^+^ decline and Ca^2+^ increase as the only rapidly LK-responding domain of the root. The positioning of the KSN confined deep inside the root tip surprises, because it reveals that plants do not recognise changes in outside K^+^ at their epidermal surface. However, this positioning facilitates multidirectional signalling to coordinate spatially separated diverse downstream responses including regulation of stem cell activity, modulation of endodermis differentiation and adjustment of gene transcription and ion transport.

KSN-born Ca^2+^ signals in turn trigger CIF-peptide signalling from the stele to enhance activity of the SGN3-LKS4/SGN1 receptor module that provokes formation of defined ROS pattern by activating distinct NOXs. Exclusive localisation of RBOHF in the Casparian strip domain (CSD) combined with the exclusion of RBOHC/D from the CSD provide a plausible separation mechanism creating further response bifurcation^22^. The resulting accelerated CS formation can therefore synergise with simultaneous transcriptome reprogramming for adjusting ion uptake and fluxes to withstand deprivation of external K^+^. This uncovers a novel concept how plants integrate environmental cues with modulation of specific developmental programmes at the molecular level.

On the organ scale, the root tip not only homes the stem cell niche (SCN) for root growth and hormonal gradients as organisers directing differentiation but also – as reported here – the KSN as organiser of organismal K^+^ homoeostasis and modulator of growth and differentiation. This sophisticated organising pattern facilitates integration of developmental programmes with K^+^ sensing to fine-tune growth and differentiation for optimising nutrient foraging.

## Methods

### Plant materials and growth conditions

The *Arabidopsis thaliana* lines used in this study, including *lks4-1* (SALK_055095), *sgn3* (SALK_064029), *cif2* (SALK_138660), *rbohC* (SALK_016593), *rbohC-1* (SALK_065153), *rbohC-2* (SALK_071801), *rbohD* (SALK_035391), *rbohD-1* (SALK_109396), *rbohD-2* (SALK_129025), *rbohB* (SAIL_749_B11), *rbohF-2* (SALK_057041), *rbohF-3* (CS9557) and *hak5* (SALK_005604) were obtained from the NASC (Nottingham Arabidopsis Stock Centre). Columbia ecotype Col-0 was used as wild type (WT). The lines pCIF1:NLS-GFP, pCIF2:NLS-GFP, pSGN1:SGN1-Cit, pSGN3:SGN3-Venus in *sgn1-2*^19,21^ were kindly provided by N. Geldner, *rbohD/RBOHD*^*S39D*^ and *rbohD/RBOHD*^*S39A*^ lines were kindly provided by J. M. Zhou^32^ and the line expressing roGFP2-Orp1^28^ was kindly provided by M. Schwarzländer.

For propagation, plants were grown in a potting soil mixture (rich soil:vermiculite, 2:1, v/v) in growth chambers (22°C, illumination 120 µmol m^-2^ s^-1^, 6 h daily light period; relative humidity 70 %).

### Generation of transgenic plants

Genomic DNA encompassing 2,515 bp including coding region and regulatory sequences of *LKS4* was cloned into pCAMBIA1300 and transformed into *lks4-1* to generate complementation lines (COM1 and COM2). The *LKS4* cDNA fused to *GFP* was cloned into pCAMBIA1391 driven by *LKS4* native promoter (2,001 bp) and transformed into *lks4-1* to obtain *lks4-1*/*ProLKS4:LKS4-GFP* lines. The *RBOHD*^*S39D*^ cDNA fused to a Flag tag together with a 2,041 bp promoter fragment was cloned into pCAMBIA1300 and transformed into *lks4-1*. T_4_ homozygous transgenic plants were used for phenotypic analyses.

Crispr/Cas9 mutagenesis was performed as described previously^33^. A pair of sgRNA targets (C1: AGTCTGAGGTCATATTTCT and C2: CGGCAACGATAGTGATATCC) *LKS4* was cloned into the pHSE401-2gR vector and used to generate *lks4-2*. The pHSE401-2gR vector containing the sgRNA targets (C1: GGTGCCTTTAGCGGTCCGCT and C2: CCAGCTGCTTGTTGGACAC) of *RBOHD* was used to generate *lks4-1 rbohD*.

The coding sequence of the ratiometric K^+^ sensor lc-LysM GEPII1.0^24^ was cloned into the pGGC vector^34^. Plant transformation constructs were generated via GreenGate assembly using the modules pUBQ10 (A), N-decoy (B), lc-LysM GEPII1.0 sequence (C), C-decoy (D), tHSP18.2 (E), pNOS-HygR-pAG7 (F) in pGGZ003. YC3.6 Ca^2+^ reporter lines were generated with the GreenGate modules pUBQ10 (A), N-decoy (B), YC3.6 (C), C-decoy (D), tHSP18.2 (E), pNOS-HygR-pAG7 (F) in pGGZ003.

All transgenic Arabidopsis thaliana lines were generated by Agrobacterium-mediated transformation using the floral dip method^35^.

### Assays for K^+^-dependent phenotyping, K^+^ content determination and NaCl tolerance

Media for phenotype analyses were prepared by modifying Murashige and Skoog (MS) medium as following: 1.5 mM MgSO_4_ remained unchanged, NH_4_NO_3_ and KNO_3_ were removed, and 1.25 mM KH_2_PO_4_ and 2.99 mM CaCl_2_ were replaced by 1.25 mM H_3_PO_4_ and 2.99 mM Ca(NO_3_)_2_, respectively. K^+^ concentrations in HK (high K^+^) and LK (low K^+^) were adjusted to 5 mM and 0.01 mM by adding KCl. Phenotype assays and K^+^ content determinations were performed as described previously^36^. For short-term LaCl_3_ treatments 50 µM LaCl_3_ was applied for 30 min and then seedlings were used for subsequent assays in the respective assay buffers. For long-term LaCl_3_ treatment 50 µM LaCl_3_ was added to the growth medium. NaCl tolerance of *rbohF* and WT was assayed as described^25^.

### RT-qPCR, transcriptomics and phospho-proteomics analyses

RT-qPCRs were performed as described previously^37^. *Actin2/8* was used as internal standard for normalisation. Three biological replicates consisting of 80-100 seedlings from two plates were used for each RT-qPCR experiment.

For transcriptomics, 5-day-old seedlings were transferred to HK or LK for 1 d. Plant roots were collected for RNAseq analyses. Paired sequence reads of 150 bp were aligned using CLC Genomics Workbench 10.0.2 (QIAGEN, Aarhus, Denmark). The Arabidopsis genome assembly of ftp.ensemblgenomes.org/pub/current/plants/fasta/arabidopsis_thaliana/dna/Arabidopsis_thaliana.TAIR10.dna.toplevel.fa.gz (downloaded 10.8.2019) was used as reference genome. The following criteria were applied to map sequence reads: mismatch cost: 2, deletion cost: 3, insertion cost: 3, length fraction: 0.8, similarity fraction: 0.8. Data were normalised by calculating transcripts per kilobase million (TPM) and annotated with Arabidopsis gene and mRNA track (ftp.ensemblgenomes.org/pub/current/plants/gtf/arabidopsis_thaliana/Arabidopsis_thaliana.TAIR10.44.gtf.gz, 10.8.2019). Differential expression analysis was performed with the Differential expression for RNA-Seq 1.0 module of CLC Genomics. Differentially expressed genes (DEGs) were defined as exhibiting ≥2-fold expression changes in the analysed condition or genotype with FDR-p≤0.05. DEGs were analysed for enriched GO terms using the PANTHER tools with default/recommended settings^38,39^. Heatmaps were drawn from TPM expression values using R (r-project.org) and the R packages ComplexHeatmap (version 2.1.0), grid and circlize (version 0.4.8)^40,41^. Venn diagrams were assembled with Create Venn Diagram for RNA-Seq 0.1 within CLC Genomics.

### BiFC and Co-immunoprecipitation assays

BiFC assays were performed as described previously^42-44^. For Co-IP, *Arabidopsis* protoplasts were transfected with the indicated plasmids and incubated overnight. Total proteins were extracted using extraction buffer (10 mM Tris-HCl [pH 7.6], 150 mM NaCl, 2 mM EDTA, 0.5% NP40, proteinase inhibitor cocktail). For anti-Myc IP, total proteins were incubated with 50 µL agarose conjugated with anti-Myc antibody (Sigma) for 2 h, and then washed for six times with washing buffer (10 mM Tris-HCl [pH 7.6], 150 mM NaCl, 2 mM EDTA, 0.5% NP40). After washing, the proteins were separated by SDS-PAGE and detected by anti-Myc and anti-Flag immunoblot.

### *In vitro* phosphorylation assays and p-site mapping

His-LKS4, GST-LKS4, GST-SGN3-C (899-1249 aa), and GST-RBOHs were expressed in *E*. *coli* and purified using Ni and glutathione sepharose beads (GE Healthcare). Site mutagenesis of RBOHC and RBOHD was performed with a QuickChange Kit (Agilent Technologies; 210518). The indicated proteins were incubated in protein kinase assay buffer (20 mM Tris-HCl [pH 7.6], 10 mM MgCl_2_, 1 mM DTT, 1 µCi [γ-^32^P] ATP and 10 µM cold ATP) at 30°C for 30 min, separated by 10% SDS-PAGE and detected on a Typhoon 9410 imager. Coomassie brilliant blue stains served as loading controls.

P-site mapping of RBOHD was initially performed by in vitro phosphorylation of N-terminal fragments (Extended Data Fig. 5b-d) that revealed fragments A and D as phosphorylated by LKS4. Subsequently, all 20 potential phosphorylation sites in these two fragments were mutated to Ala (A) one by one for target site identification (Extended Data Fig 5e-h).

### ROS staining and ratiometric ROS imaging

Seedlings grown on HK for 5 d were transferred to LK or HK for 24 h. ROS staining of seedlings was performed by incubation with 10 µM H_2_DCFDA (2,7-dichlorodihydrofluorescein diacetate, SIGMA) for 2 min. Fluorescence was observed using a confocal laser scanning microscope (LSM880, Carl Zeiss; excitation at 488 nm, emission at 490-560 nm).

For ratiometric ROS imaging, seedlings of roGFP2-Orp1 were grown for 4 d on HK plates and transferred to HK or LK liquid medium for 24 h. roGFP imaging was performed using a TCS SP8 X confocal laser scanning microscope. roGFP-Orp1 was sequentially excited at 405 nm and 488 nm and GFP emission was detected at 500 - 535 nm with the pinhole set to 5 airy units. Background fluorescence with excitation at 405 nm and collected at 425 - 475 nm^28^. Background fluorescence was subtracted, and 405/488 ratios were calculated with ImageJ (imagej.nih.gov) software with the RatioPlus plug-in.

### PI stains, CS formation and suberisation assays

CS formation assays were performed double-blinded in parallel in Münster and Beijing. Seedlings were grown on HK for 4-5 d, and transferred to LK or HK for 24 h/48 h. Seedlings were incubated with 15 µM PI (Propidium iodide, SIGMA) for 10 min. Fluorescence was observed with a 20x objective using confocal laser scanning microscopes (LSM880, Carl Zeiss (Beijing) or Leica TCS SP8 X (Münster); excitation at 561 nm, emission at 580-640 nm or 600-700 nm). To assay CS formation, cells in the cortex cell file were counted from the first cell that was at least twice as long as wide until the staining of the stele ended. Suberisation was determined as described previously^13^.

### Luminescence observation, ratiometric K^+^ and Ca^^2^+^ imaging

*ProHAK5:LUC* plants were grown on HK for 5 d and transferred to LK or HK for 1 d. Seedlings were sprayed with luciferin solution (0.5 mg/mL) and kept in darkness for 20 min before luminescence was monitored using a Lumazone CCD camera (CA1300B, Roper Scientific) and quantified using WinView32 software.

Ratiometric K^+^ imaging was performed as described previously with the changes that plants expressing lc-LysM GEPII1.0 (that combines K^+^-binding domains of the bacterial K^+^ binding protein (KBP) with CFP and YFP variants to enable FRET-based recordings and has a K_d_ for K^+^ of 27 mM) instead of YC3.6 were investigated and that the incubation buffer (IB) contained 10 mM KCl instead of 5 mM KCl^11^. For long-term K^+^ deprivation, seedlings were grown for 4 d on HK, transferred to petri dishes with LK or HK and incubated for up to 48 h. Subsequently, seedlings were mounted on an object slide for microscopic analyses (performed either on a Zeiss microscope as described in^45^ or on a Leica SP5 microscope as described in^24^). FRET efficiency values indicative for K^+^ accumulation were determined as ratio value (R) obtained by dividing the normalised emission intensities of YFP and CFP at defined timepoints (ΔR/R_0_) as described previously for the Ca^2+^ sensor YC3.6^45^. Ca^2+^ imaging was performed as described previously^11^.

### Longitudinal quantification of reporter gene expression and statistical analyses

Plants were grown for 3-4 d on HK and then transferred to LK for up to 24 h. GFP fluorescence was recorded with a Zeiss microscope using a 10x objective. Single stage images were stitched using the ImageJ Plugin Pairwise stitching to obtain composed images. Background fluorescence was subtracted. A longitudinal line-scan ranging from the QC to the mature DZ was performed along the stele and intensity plotting was performed using ImageJ. False color representations were generated with the orange-icb LUT. Statistical analyses were performed within Sigmaplot 12.5 (Systat Software, Erkrath, Germany) or GraphPad Prism 8.2.1 (GraphPad Software, San Diego, CA, USA). For multiple comparison analyses appropriate test were chosen according to data distributions.

### Accession numbers

Sequence data from this article can be found in the *Arabidopsis* Genome Initiative or GenBank/EMBL databases under the following accession numbers: *LKS4* (*AT1G61590*), *SGN3* (*AT4G20140*), *CIF2* (*AT4G34600*), *RBOHB* (*AT1G09090*), *RBOHC* (*AT5G51060*), *RBOHD* (*AT5G47910*), *RBOHF* (*AT1G64060*), *HAK5* (*AT4G13420*), *BAK1* (*AT4G33430*).

## Acknowledgements

We are gratefully indebted to N. Geldner, S. Fujita and J. M. Zhou for helpful discussion and providing various materials. We are grateful to M. Schwarzländer for providing roGFP-Orp1 lines and mammalian GEPII plasmids. We thank Q. J. Chen for providing Crispr/Cas9 vectors. We gratefully acknowledge support from J. B. Kroll and A. K. van Wüllen in CIF2 expression data analyses, P. Manishankar for sharing data on *rboh* mutant crosses and thank K. H. Edel for helpful discussions. This work was supported by the Chinese Universities Scientific Fund (2019TC122 and 2019TC228 awarded to Y.W.), a grant from the National Natural Science Foundation of China (NSFC, 31921001 to Y.W.) and a Joint Sino-German Research Project financed by the NSFC (31761133011 to Y.W.) and the Deutsche Forschungsgemeinschaft (DFG, 391703796 to J.K.). G.H. was supported by a fellowship from the Chinese Scholarship Council (CSC, 201806350012).

## Author contributions

F.L.W., Y.L.T., L.W., X.Q.D., A.E., Z.F.W., G.F.H., J.P.H. and I.S.T performed experiments and analysed data. F.L.W., L.W., J.K. and Y.W. wrote the manuscript with input from all authors. F.L.W., Y.L.T., L.W., and A.E. assembled the figures. J.K. and Y.W. conceived and J.K., Y.W. and W.H.W supervised the project.

## Competing interests

The authors declare no competing interests.

**Extended Data Figure 1.**
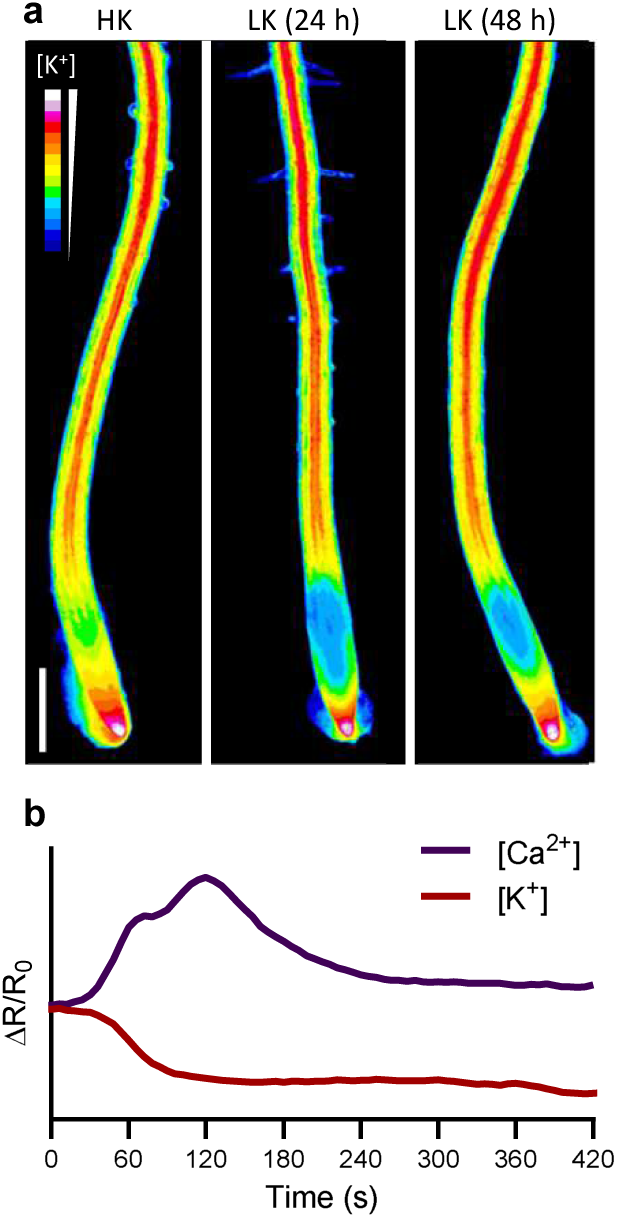
Long-term K^+^ imaging reveals differential dynamics of cytoplasmic K^+^ concentration in specific cell types and root zones. **a**, False color representation of cytoplasmic K^+^ concentration after exposure to LK for 24 h and 48 h. Scale bar, 200 µm. **b**, K^+^ and Ca^2+^ dynamics in the EZ after onset of K^+^ depletion. The depicted graphs represent real experimentally obtained values (K^+^, n=6; Ca^2+^, n=4) and only for optical clarity, error bars and the preincubation period of the experiments were omitted.

**Extended Data Figure 2.**
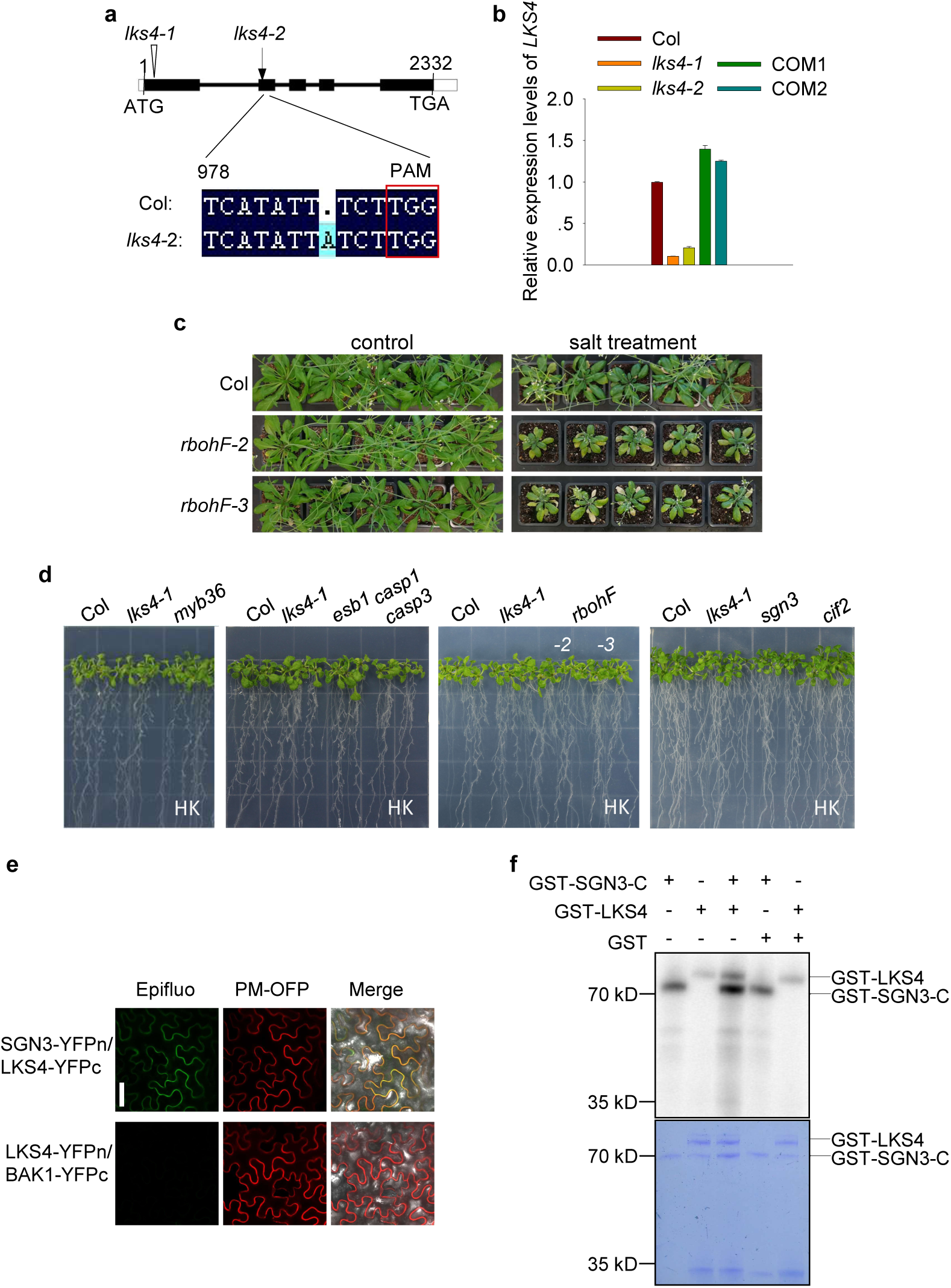
Characterisation of LKS4, *lks4* alleles and CS-related mutants. **a**, Presentation of LKS4 gene structure and the characterized mutant alleles. Filled boxes and lines represent exons and introns. The T-DNA insertion site in *lks4-1* and the mutation in *lks4-2* are indicated by triangle and arrow. Sequence details of the single nucleotide insertion in *lks4-2* in comparison to the corresponding Col (WT) sequence. **b**, RT-qPCR analyses of *LKS4* expression in the indicated genotypes. **c**, Salt stress phenotypes of different *rbohF* alleles grown in soil. **d**, Phenotypes of the indicated genotypes grown on HK. **e**, BiFC assays combining LKS4 with SGN3 or BAK1. PM, plasma membrane; Scale bar, 50 µm. **f**, Autoradiograph and Coomassie gel of *in vitro* phosphorylation assays combining LKS4 with the kinase domain of SGN3 (SGN3-C).

**Extended Data Table 1.**
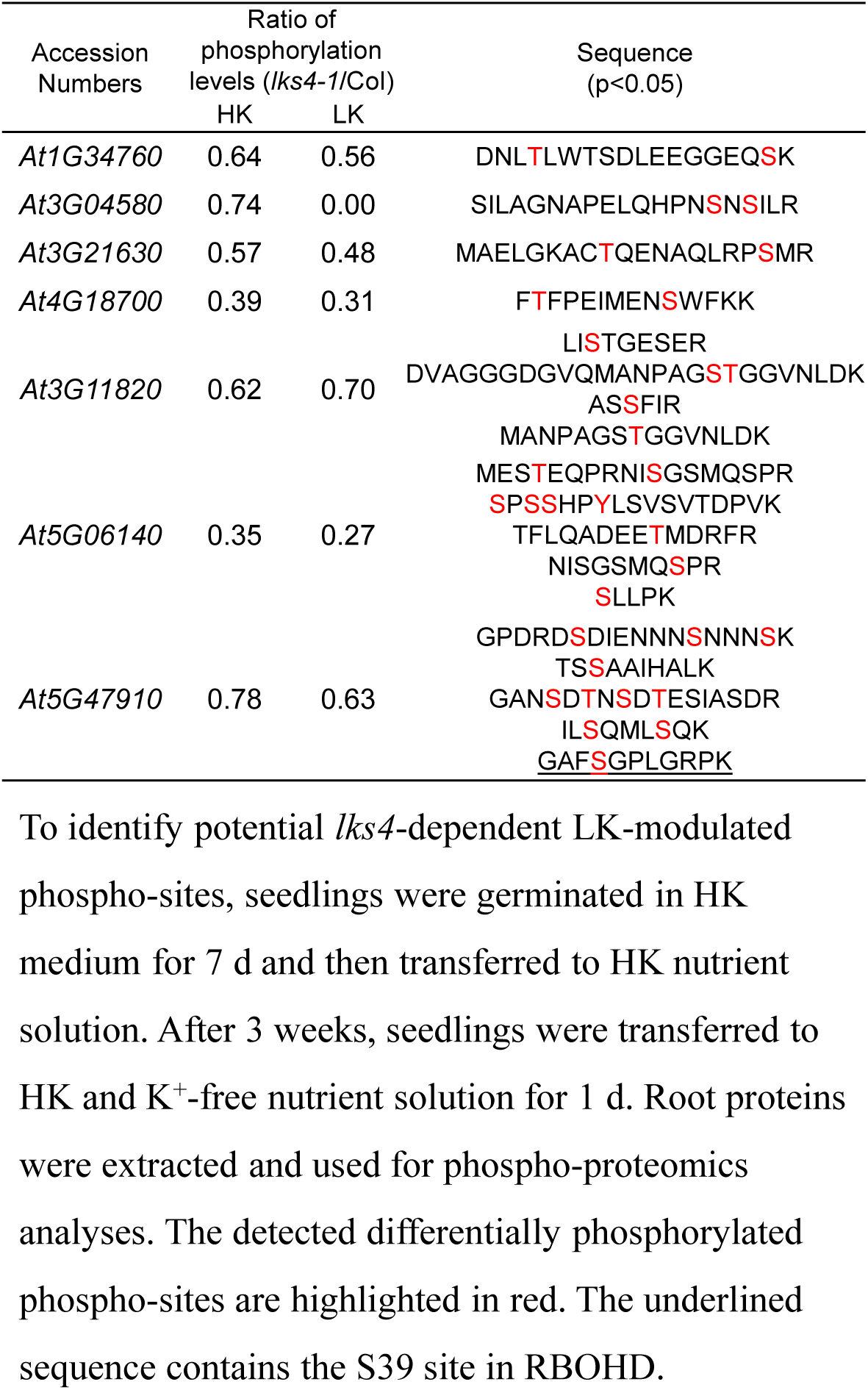
**Proteins with reduced phosphorylation in *lks4-1***. To identify potential *lks4*-dependent LK-modulated phospho-sites, seedlings were germinated in HK medium for 7 d and then transferred to HK nutrient solution. After 3 weeks, seedlings were transferred to HK and K^+^-free nutrient solution for 1 d. Root proteins were extracted and used for phospho-proteomics analyses. The detected differentially phosphorylated phospho-sites are highlighted in red. The underlined sequence contains the S39 site in RBOHD.

**Extended Data Figure 3.**
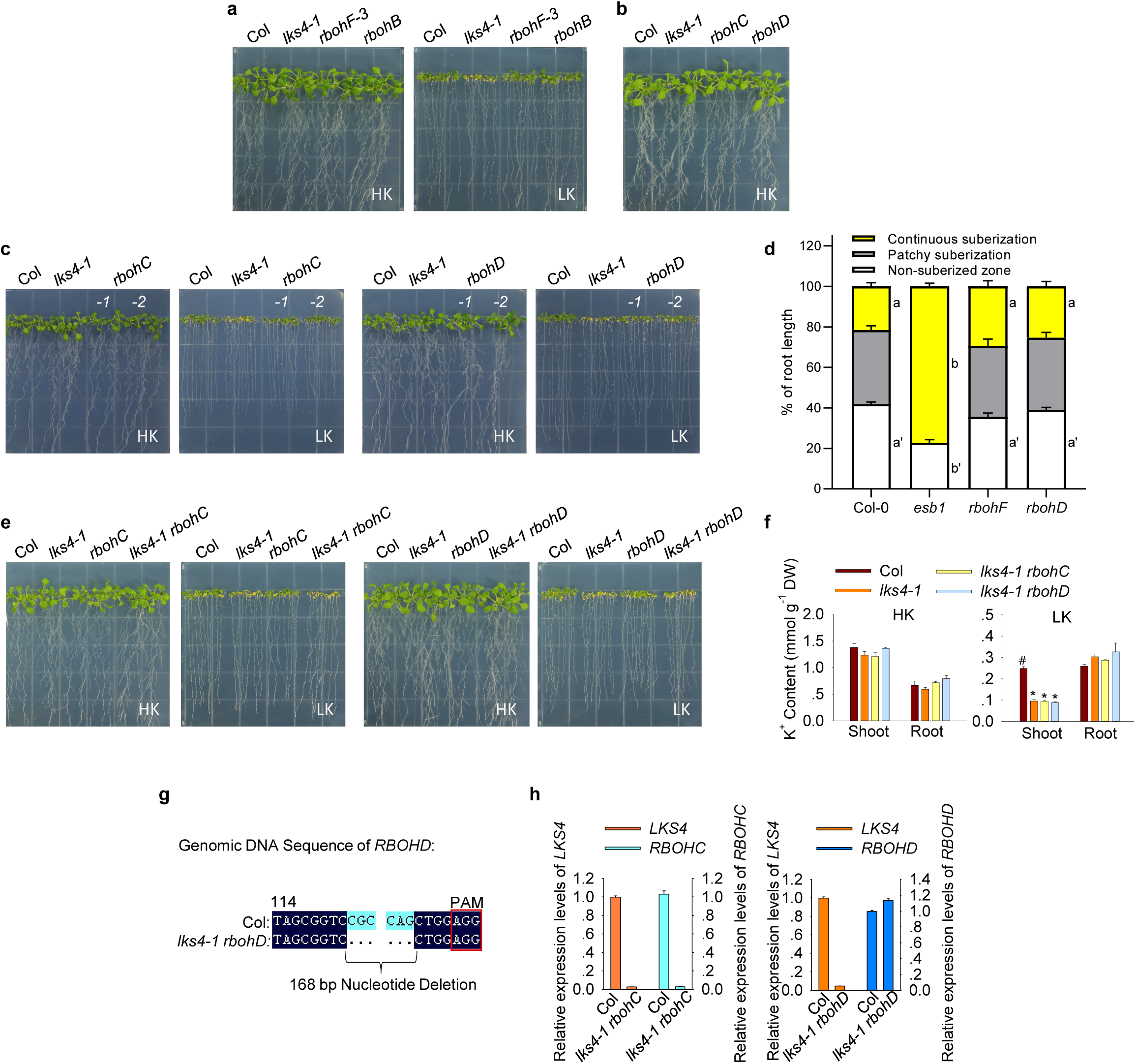
Characterisation of root-expressed NADPH oxidases. **a-c** and **e**, Phenotypes of the indicated genotypes grown on HK and LK. **d**, Analyses of CS suberisation of the indicated genotypes. **f**, K^+^ content of indicated genotypes. Mean±SE; n=3; DW, dry weight; #, control; *p<0.05. **g**, Sequence details of the 168 bp nucleotide deletion in the Crispr/Cas9-induced *rbohD* mutant allele generated in the *lks4-1* background. **h**, Verification of *LKS4, RBOHC* and *RBOHD* expression in the indicated plant lines by RT-qPCR.

**Extended Data Figure 4.**
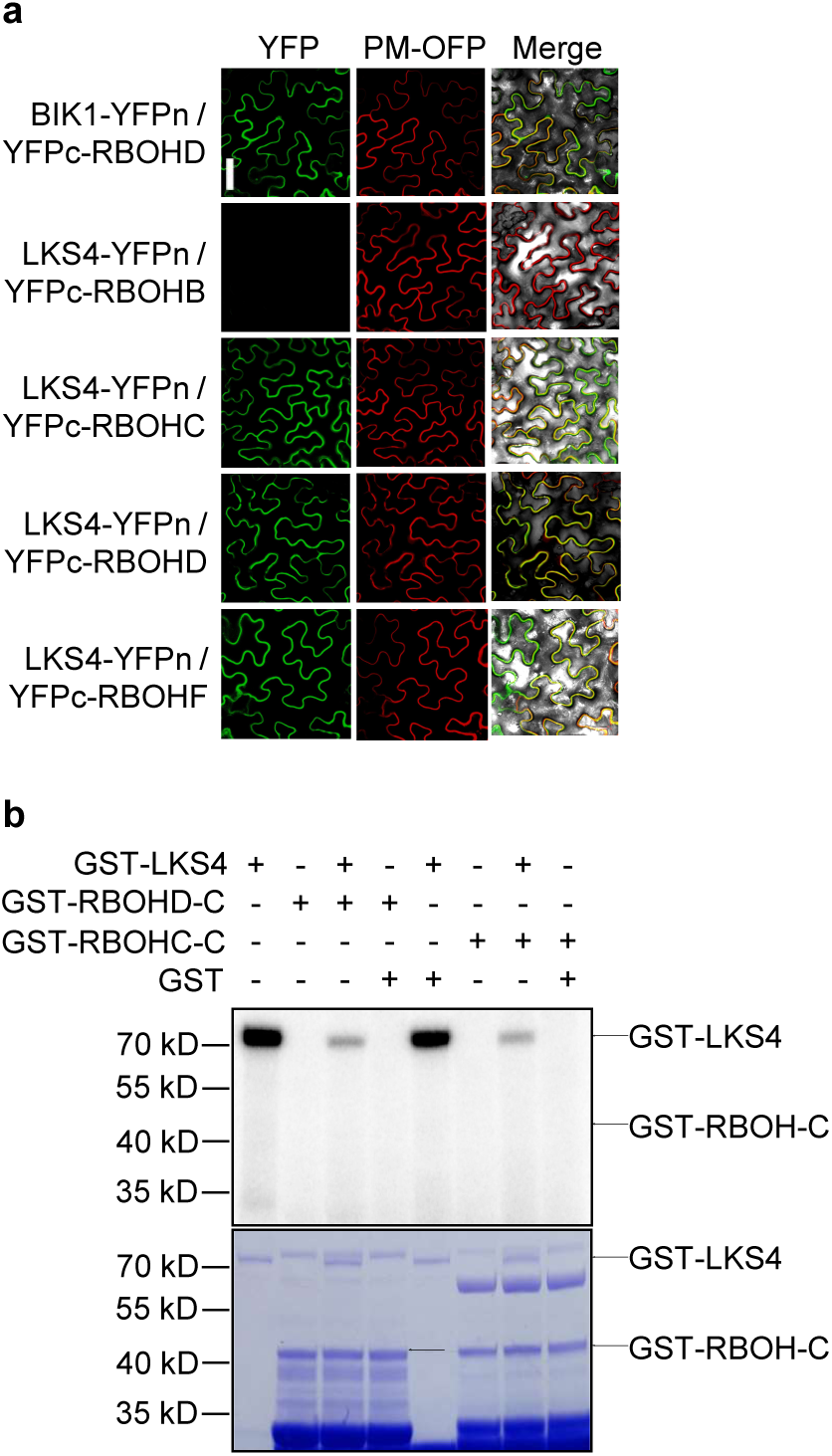
CD-phosphor. **a**, BiFC assays combining LKS4 with different RBOH proteins. CBL1n-OFP was used as a PM marker^46^ Scale bar, 50 µm. **b**, Autoradiograph and Coomassie gel of *in vitro* phosphorylation assays combining LKS4 with the C-termini of RBOHD or RBOHC.

**Extended Data Figure 5.**
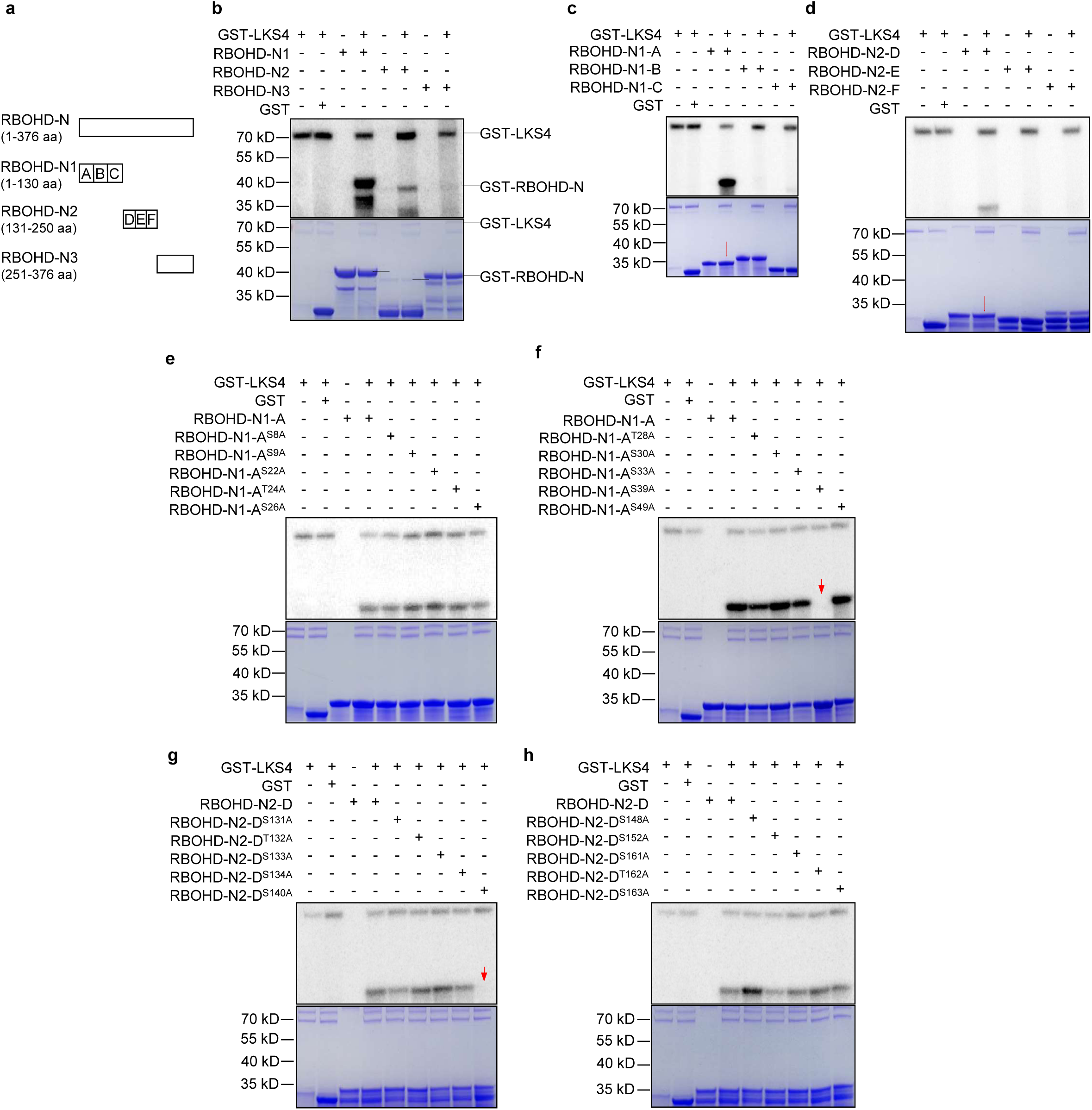
Mapping of RBOHD phospho-sites. **a**, Schematic depiction of the analysed sub-fragments. **b-d**, Mapping of phosphorylated sub-fragments of the N-terminal cytoplasmic domain of RBOHD. **b**, Phosphorylation assays comprising fragments N1, N2 and N3. **c**, Phosphorylation assays comprising fragments A, B and C. **d**, Phosphorylation assays comprising fragments D, E and F. **e-h**, Mapping of phosphorylation sites within the sub-fragments N1 and N2 of the N terminal cytoplasmic domain of RBOHD. **e**, Phosphorylation assays for analysing the residues S8, S9, S22, T24 and S26. **f**, Phosphorylation assays for analysing the residues T28, S30, S33, S39 and S49. **g**, Phosphorylation assays for analysing the residues S131, T132, S133, S134 and S140. **h**, Phosphorylation assays for analysing the residues S148, S152, S161, T162 and S163.

**Extended Data Figure 6.**
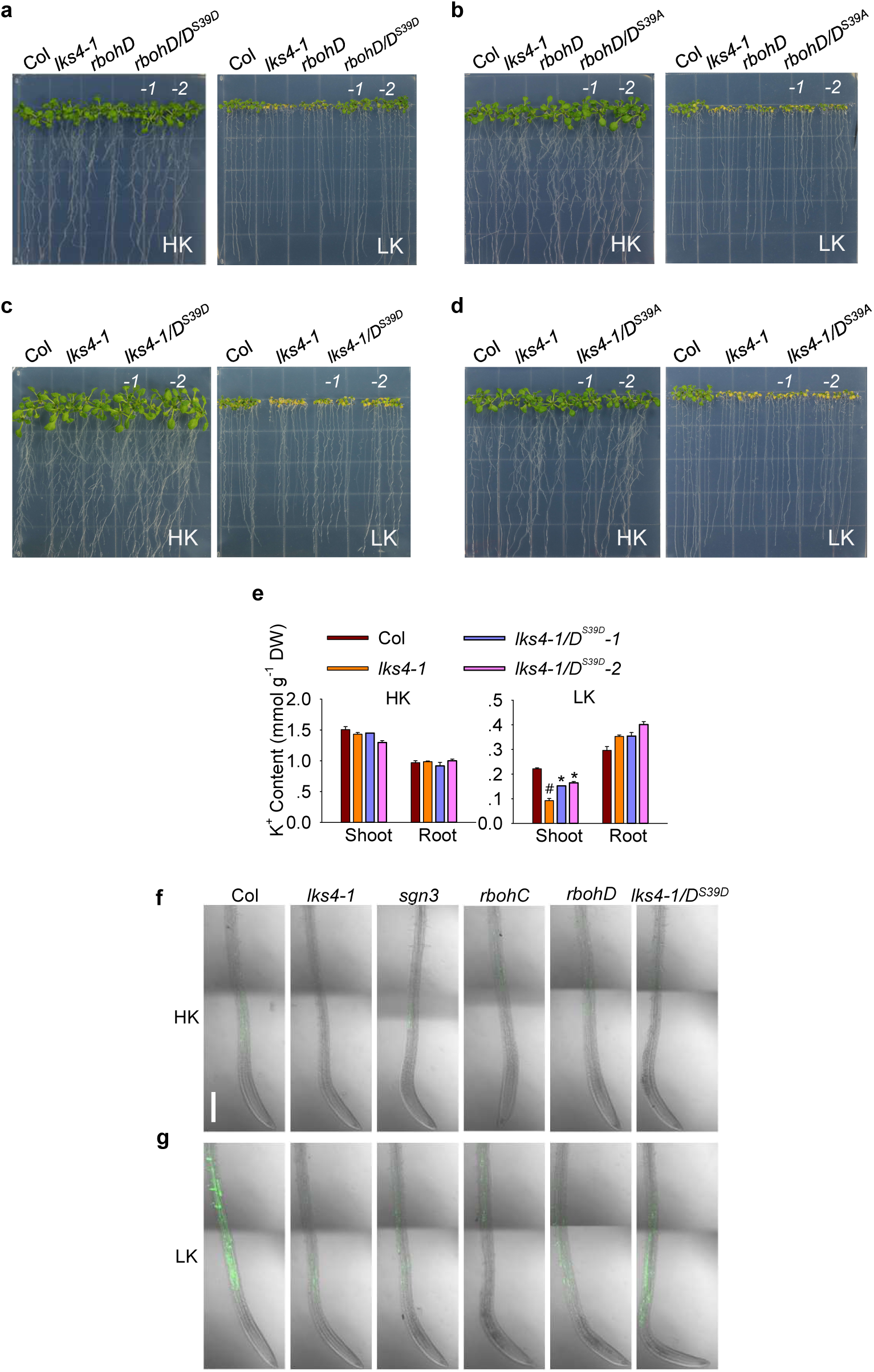
Analysis of RBOHD S39 phosphorylation and ROS pattern in mutant plants. **a-d**, Phenotypes of the indicated genotypes grown on HK and LK. **e**, K^+^ content of plants expressing RBOHD^S39D^ (D^S39D^) in *lks4-1*. Mean±SE; n=3; DW, dry weight; #, control; *p<0.05. **f**, ROS accumulation of indicated genotypes after 24 h LK exposure reported by H_2_DCFDA fluorescence. Scale bar, 200 µm.

**Extended Data Figure 7.**
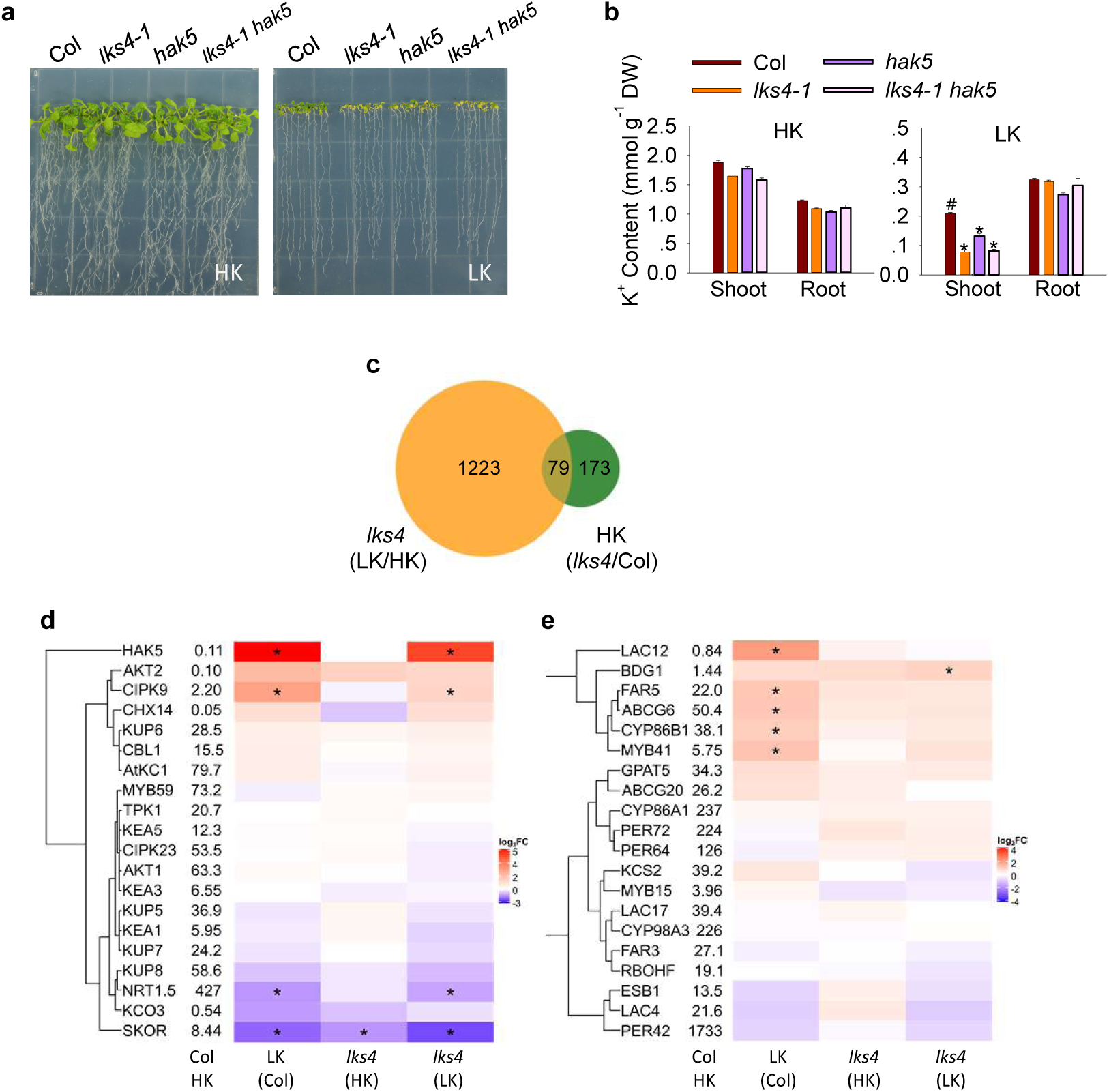
Transcriptome analyses for characterising the impact of K^+^ deprivation in Col and *lks4*. **a** and **b**, Phenotypes and K^+^ content of indicated genotypes grown on HK and LK. Mean±SE; n=3; DW, dry weight; #, control; *p<0.05. **c**, Venn diagram illustrating differentially expressed genes (DEGs) in Col (WT) and *lks4* upon exposure to HK or LK. **d** and **e**, Heatmap illustrating expression changes of genes related to K^+^ homoeostasis (d) and suberin and lignin (e). *DEGs with at least 2-fold expression change.

**Extended Data Table 2.**
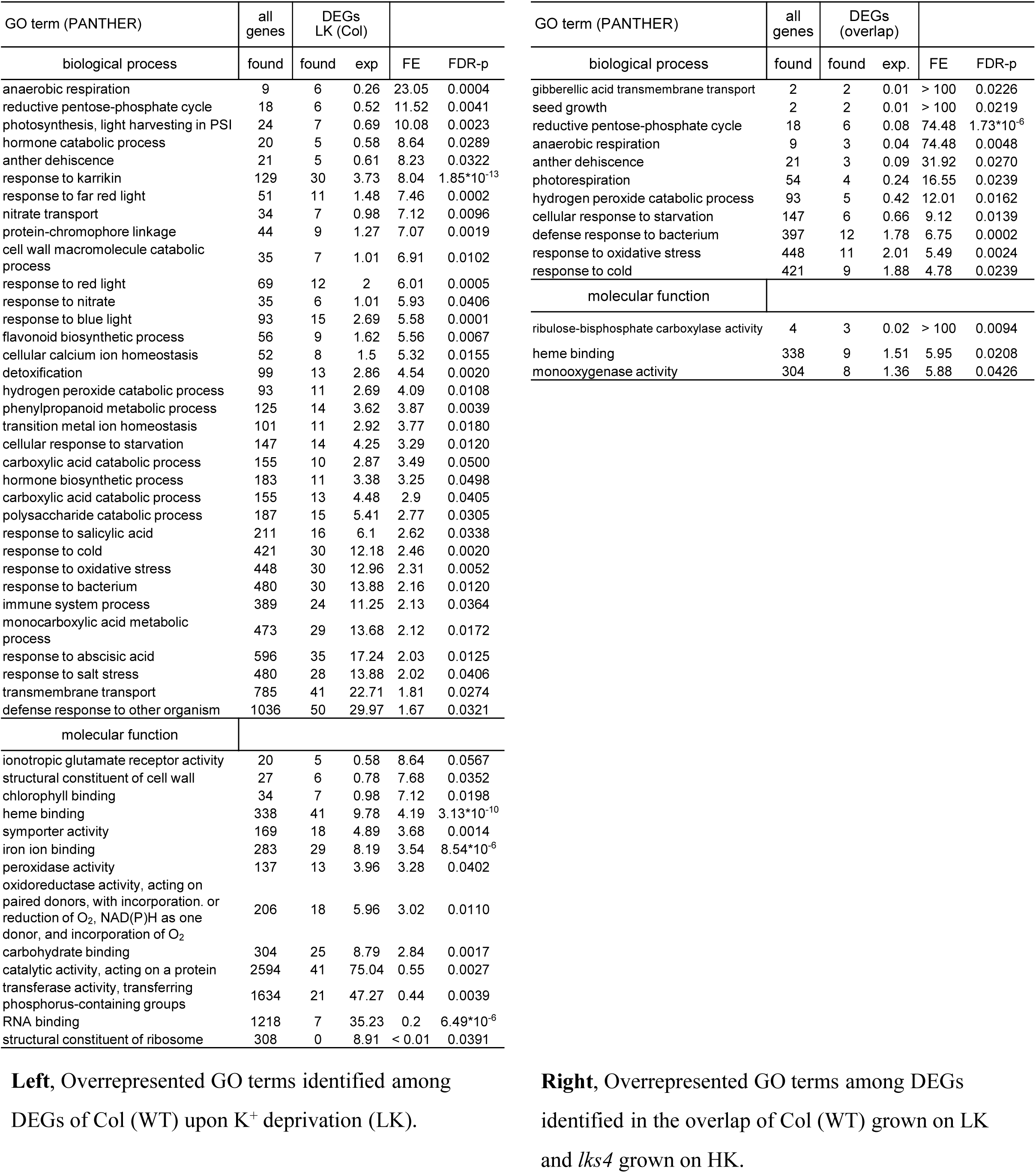
**Enriched GO terms**. Left, Overrepresented GO terms identified among DEGs of Col (WT) upon K^+^ deprivation (LK). **Right**, Overrepresented GO terms among DEGs identified in the overlap of Col (WT) grown on LK and *lks4* grown on HK.

**Extended Data Figure 8.**
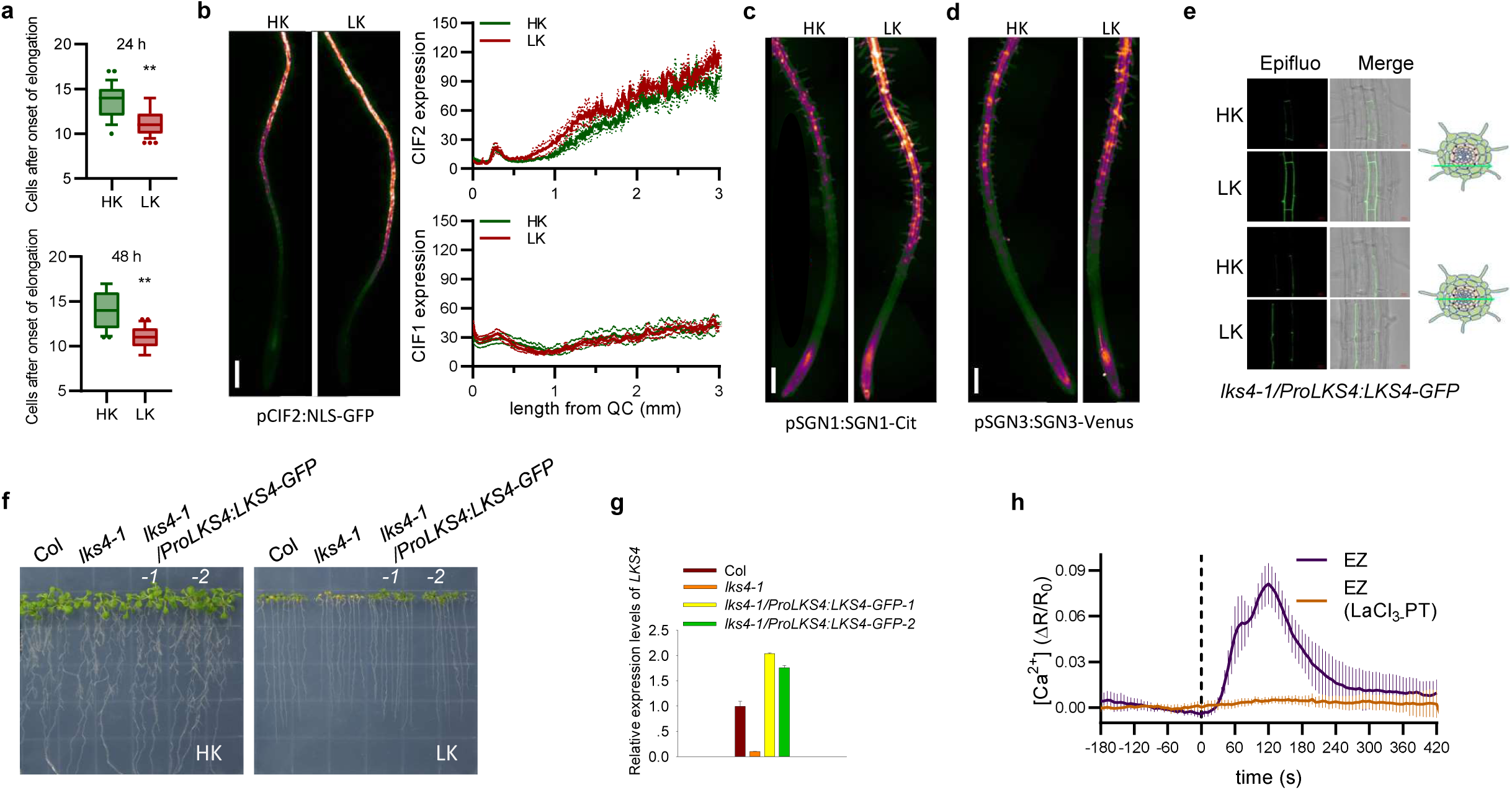
LK-induced Ca2^+^ signalling accelerates CS formation and plant resilience against K^+^ deprivation. **a**, Cortex cell number from the onset of elongation until PI staining in the stele ends after 24 h and 48 h HK or LK exposure. Median and 10-90 percentile; n=27, 34, 20, 21; HK 24 h, LK 24 h, HK 48 h, LK 48 h from 4 independent experiments; **p<0.01. **b**, Left, CIF2 expression displayed as false colour GFP intensity of pCIF2:NLS-GFP expressing seedlings exposed to HK or LK for 6 h. Scale bar, 200 µm. Right upper panel, Longitudinal quantification of *CIF2* expression represented as 8-bit GFP intensity. HK or LK treatment for 6 h. Mean±SEM; n=3, 4; HK, LK. Right lower panel, Quantification of CIF1 expression represented as 8-bit GFP intensity. Mean±SEM; without background subtraction; n=5, 6; HK, LK. **c**, LKS4/SGN1 expression displayed as false colour GFP intensity of pSGN1:SGN1-Cit plants exposed to HK or LK for 24 h. Scale bar, 200 µm. **d**, SGN3 expression displayed as false colour GFP intensity of pSGN3:SGN3-Venus in *sgn1-2* exposed to HK or LK for 24 h. Scale bar, 200 µm. **e**, GFP fluorescence observation showing the protein expression of LKS4 under LK and HK conditions. Seedlings of *lks4- 1*/*ProLKS4:LKS4-GFP* complementation lines were grown on LK and HK medium for 5 d. Green arrows in the cross sections of roots indicate observation positions. **f** and **g**, Phenotypes of and *LKS4* expression (analysed by RT-qPCR) in *lks4-1*/*ProLKS4:LKS4-GFP* complementation lines grown on HK and LK for 12 d. **h**, Quantification of the primary Ca^2+^ signal upon K^+^ depletion (starting at 0 s) displayed as normalised ratio changes. Mean±SEM; n=4; La^3+^-PT, incubation with 50 µM LaCl_3_ prior to Ca^2+^ imaging (without LaC_3_).

**Supplementary Table 1.**
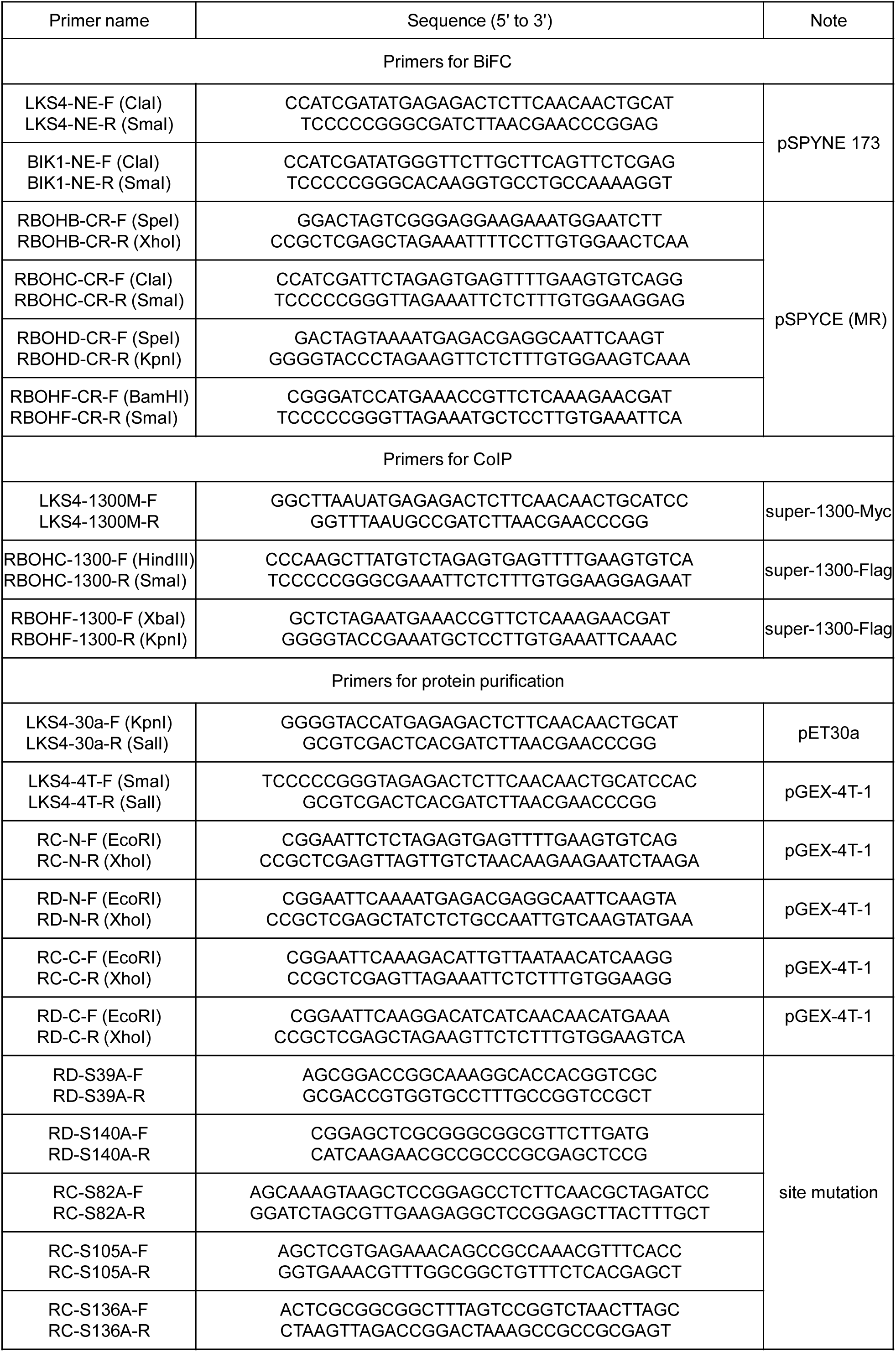
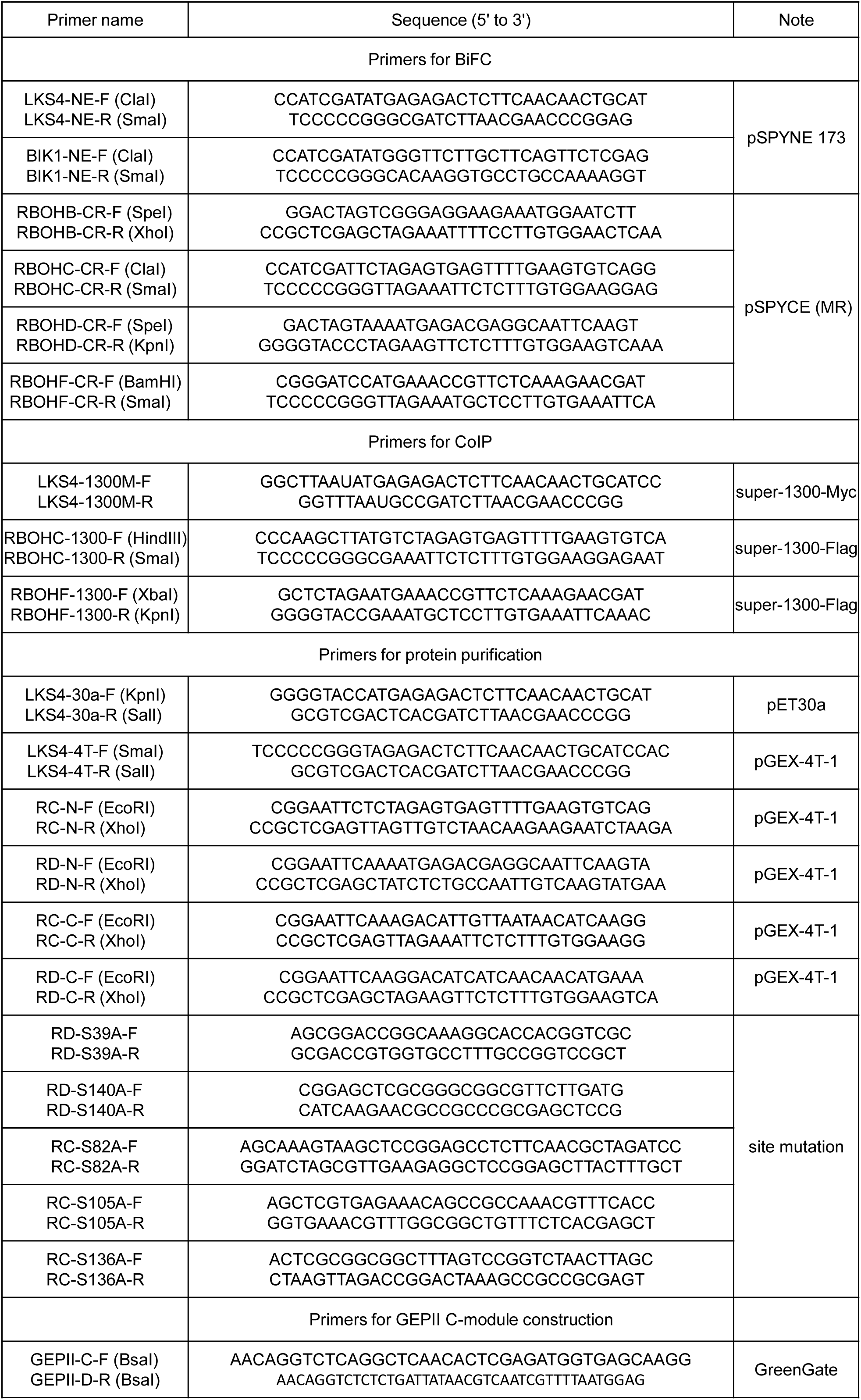
**Sequences of Primers used in this study**.

## Supplementary information

**Supplementary Table 1** | **Sequences of primers used in this study**.

**Supplementary Table 2** | **Transcriptomics analyses of K**^**+**^ **availability-modulated gene expression in WT and *lks4-1***. List of significantly differentially expressed genes (at least 2-fold change in expression, FDR-p<0.05) in roots of Col (WT) and *lks4-1* seedlings exposed to HK or LK for 24 h, respectively. **a** and **b**, up- and downregulated genes in Col (LK/HK). **c** and **d**, up- and downregulated genes in *lks4-1* (LK/HK). **e** and **f**, up- and downregulated genes on HK (*lks4-1*/Col). **g** and **h**, up- and downregulated genes on LK (*lks4-1*/Col)

**Supplementary Video 1** | **Live imaging of cytosolic K**^**+**^ **dynamics of an Arabidopsis root upon K**^**+**^ **depletion in the media**. Time-lapse imaging of cytosolic K^+^, represented in false colour. IB, incubation buffer; 0K+, 0K^+^ buffer.

## Data availability

Accession numbers of all the Arabidopsis genes analysed in this study are listed in Methods. All RNA-seq data from this study will be deposited on a public archive server before publication. Seeds of transgenic lines used in this study are available from the corresponding authors upon reasonable request.

